# To group or not to group: group size dynamics and intestinal parasites in Indian peafowl populations

**DOI:** 10.1101/2020.01.03.893503

**Authors:** Priyanka Dange, Pranav Mhaisalkar, Dhanashree Paranjpe

## Abstract

Animals can form groups for various reasons including safety from predators, access to potential mates and benefits of allo-parental care. However, there are costs associated with living in a group such as competition for food and/or mates with other members of the group, higher chances of disease transmission, etc. Group size dynamics can change with the biotic and abiotic environment around individuals. In the current study, we explored the links between group size dynamics and intestinal parasites of Indian peafowl (*Pavo cristatus*) in the context of seasons and food provisioning. Data for group size was collected across three seasons (Pre-Monsoon, Monsoon and Post-Monsoon) at three field sites (Morachi Chincholi, Nashik and Rajasthan). Individual and group sightings of peafowl were noted down along with group size and composition (no. of males, females, adults, juveniles, sub-adults). Faecal samples were collected from food provision and non-provision areas across the same three seasons at same field sites. Parasite load in the samples was quantified using microscopic examination. Group size was significantly higher in Pre-Monsoon season as compared to Monsoon and Post-Monsoon seasons. Monsoon and Post-Monsoon seasons had higher intestinal parasite prevalence and load as compared to Pre-Monsoon season. Thus, group size and intestinal parasites of Indian peafowl have an inverse relationship across seasons. Parasite load was significantly greater at food provision sites as compared to non-provision sites while parasite prevalence was comparable. Aggregation of individuals at the food provision sites may influence the parasite transmission and group-size dynamics in Indian peafowl. In conclusion, Indian peafowl are behaviourally plastic and fission-fusion of social groups may allow them to tackle ecological pressures such as predation and parasite transmission in different seasons.

## Introduction

Animals can form groups for various reasons that include safety from predators (Powell 1974; Pulliam 1973; Silk 2007), finding food (Clark and Mangel 1986; Silk 2007), breeding (Silk 2007), and benefits of allo-parental care (Wittemeyer et al. 2005). However, there are also costs associated with living in a group such as competition for food (Krause and Ruxton, 2002) and/or mates (Silk 2007) with other members of the group, higher chances of disease transmission (Hochberg 1991), more negative interactions such as harassment (Clutton-Brock and Parker 1995), aggression (Krause and Ruxton 2002), and becoming more conspicuous to predators (Cresswell 1994; Krause and Ruxton 2002). Social groups form when the benefits of being in a group outweigh the costs (Sutton 2019). These costs and benefits can also be dynamic depending on environmental factors such as resource availability (Sutton 2019; Wittemeyer et al. 2005) and predation risk (Cresswell 1994; Krause and Ruxton, 2002). Ultimately these cost and benefits may influence the decision to be part of the group or not.

In many group-living species, groups can split (fission) or merge (fusion) as they move through the environment (Couzin 2006). Fission-fusion societies are thought to be good at adjusting to rapid changes in local environment and balancing the cost-benefits of grouping by changing the group sizes (Aureli et al. 2008). Fission-fusion societies have been mostly reported in long-lived and cognitively complex organisms such as dolphins (Smith et al. 2016), giraffe (Sutton 2019), chimpanzees (Lehmann and Boesch 2004), orangutans (van Schaik 1999) and elephants (Fishlock et al. 2013; Nandini et al. 2017). These types of grouping structures have also been reported in bats (Popa-Lisseanu 2008; Kashima et al. 2013), guppies (Croft et al. 2004) and avian systems (Silk et al. 2014). In shorebirds and wildfowl, lower amount of food resources and changes in predation risk during tidal cycles influenced their grouping decisions and consequently, fission-fusion dynamics (Fleischer 1983; Inger et al. 2006; Beauchamp 2010).

Fission-fusion dynamics are also known to influence endo-parasite transmission and susceptibility in brown-spider monkeys and gorillas (Caillaud et al. 2006; Rimbach et al. 2015). Flocking behaviour affected *Plasmodium* and *Haemoproteus* infections in Afro-tropical birds (Lutz et al. 2015). Parasite infections were also reported to change with respect to season in backyard poultry (Bhatt et al. 2014) and communal goats of Zimbabwe (Pandey et al. 1994).

Indian peafowl (*Pavo cristatus*) is a species that has co-habited human dominated landscapes for centuries in its native geographical range. This avian species is native to the Indian subcontinent and has been introduced in many parts of the world relatively recently. Although the species’ native habitat is undergrowth in open forest and woodlands near a waterbody, it is also known to occur near farmlands, villages and increasingly becoming common in urban and semi-urban areas (Burton and Burton 2002). Indian peafowls spend much of their active time on the ground. Social organization of feral Indian peafowl has been studied in context of mating behaviour (Rand et al. 1984), but it is not known if their grouping behaviour changes across space and time.

Previous study by Paranjpe and Dange (2020) showed that food provisioning changes feeding behaviours of Indian peafowl. Provisioning is known to affect parasite infections in racoons (Gompper and Wright 2005) and elk populations (Cross et al. 2007). Supplementary food available at bird feeders is known to have far reaching consequences on avian ecology in terms of increasing survival during overwintering, enhanced breeding success, changing sex-ratios of offspring in smaller avian species and range expansion of species (Robb et al. 2008). It is not known if provisioning can affect grouping behaviour and parasite infections in Indian peafowl populations. Therefore in this study, we tried to explore the links between the intestinal parasite infections and group dynamics in Indian peafowl populations in context of seasonal gradients and food provisioning (Fig 1).

**Fig 1.**
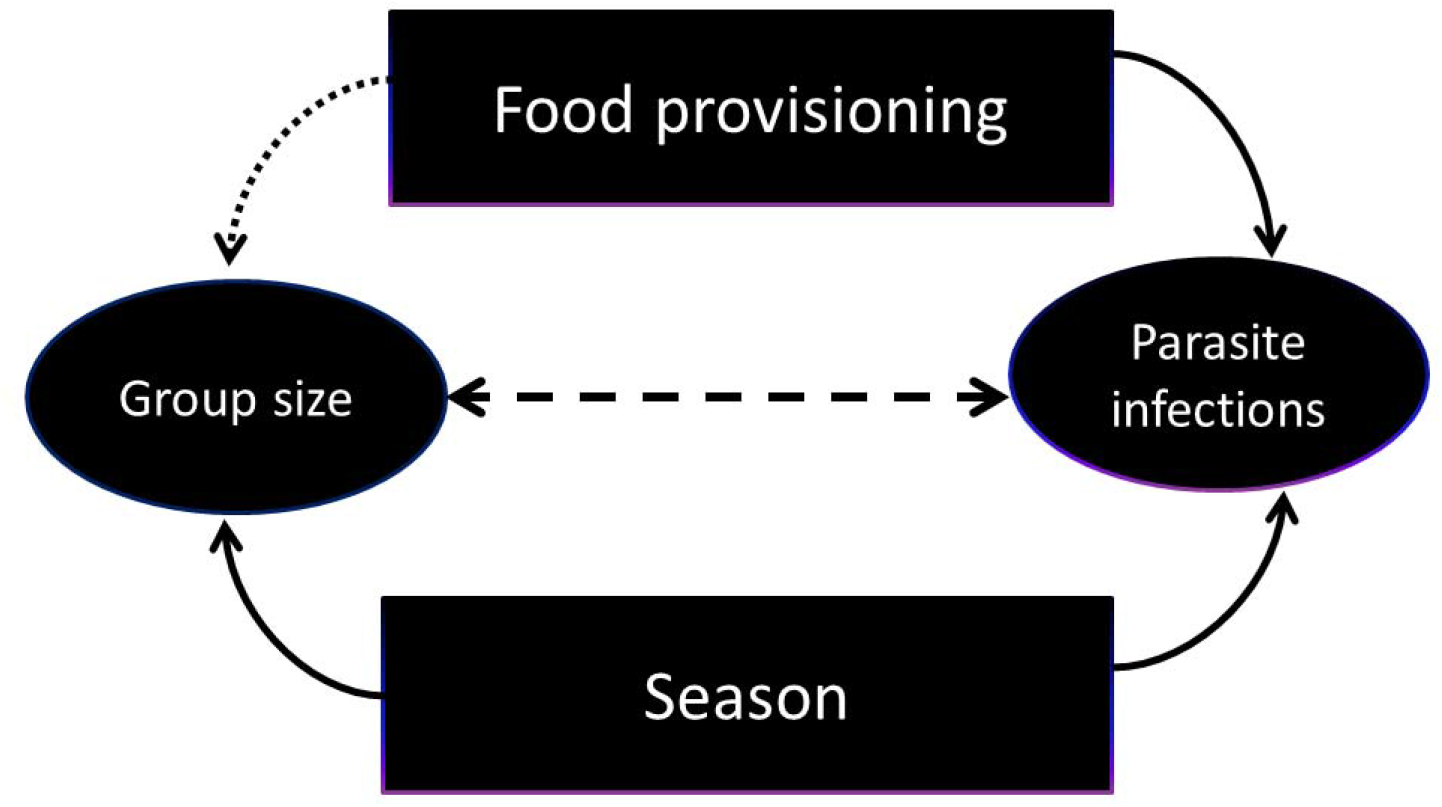
Conceptual framework of the study. This study explores the links between intestinal parasite infections and group size dynamics in Indian peafowl populations. Seasons (Pre-Monsoon, Monsoon, Post-Monsoon) may directly influence group size and intestinal parasite infections. We checked whether parasite prevalence and load is influenced by food provisioning. We speculate that food provisioning may indirectly influence group size dynamics of Indian peafowl

## Materials and methods

### Study sites

This study was conducted at the following field sites: Morachi Chincholi, Nashik and Rajasthan (villages on the periphery of Ranthambhore Tiger Reserve) from 2016 to 2019 (Table 1).

**Table 1.** Brief description of field sites.

| **Field sites** | **GPS co-ordinates and elevation** | **Provisioning of food grains (in kg) per day** |
| --- | --- | --- |
| Morachi Chincholi | 18° 48' 26.9" N, 74° 09' 29.3" E,<br>elevation 641m | ~30-35 |
| Villages in the vicinity of<br>Ranthambhore Tiger Reserve |  |  |
| 1) Govindpura | 26° 04.826" N, 76° 48.097" E,<br>elevation 227m | ~15 |
| 2) Bangarda Kalan | 26°03'57.78"N, 76°42'35.76"E,<br>elevation 225m |  |
| 3) Kuttalpur | 26° 03' 58.5" N, 76° 26' 13.4" E,<br>elevation 261m |  |
| 4) Shyampur | 26 01' 31.6" N, 76 23' 20.5" E<br>elevation 265m |  |
| 5) Sanwata-Kalakhora | 26°06'41.46"N, 76°38'33.81"E,<br>elevation 267m |  |
| Nashik | 20° 01'11.0"N, 73° 48'04.0"E,<br>elevation 404m | ~ 3-5 |

Selection of study areas was based on several criteria such as their proximity to human habitation, accessibility to the field sites throughout the year and history of peafowl populations in the area. Areas located outside protected areas and close to human habitation were given preference since food provisioning is likely to happen in these areas and studying the effects of food provisioning on grouping and parasite infections was one of the objectives of the study.

According to our estimate, ∼30-35 Kg food grains are offered to peafowl per day in the village and surrounding areas of Morachi Chincholi, while ∼15Kg grains are offered per day around homes and temple premises in the villages in Rajasthan included in this study. In both these study areas, grains are offered at designated places everyday throughout the year. Thus, Indian peafowl population in Morachi Chincholi had access to diet rich in carbohydrates (cereals) and proteins (pulses) throughout the year (Paranjpe and Dange, 2020). As much as 71% of their diet consisted of food provisioned indirectly in the form of crops or directly as grains offered in the village surroundings. Villages on the periphery of Ranthambhore Tiger Reserve (RTR), in Rajasthan offer relatively less variety of cereals and very few pulses, yet the diet of peafowl in Rajasthan has up to 61% of food provisioned by humans versus 39% natural food. The peafowl population at the third field site in Nashik, on the other hand, has access to less reliable and less varied food offered by humans (∼ 3-5 Kg per day). As a result only about 40% of their diet consists of provisioned food (Paranjpe and Dange, 2020).

### Group size

Populations of Indian peafowl were studied throughout the year using direct field observations during morning (6 am to 10 am) and evening (4 pm to 7 pm) at selected field sites. Individual and group sightings of peafowl were noted down along with date, time of day, habitat, group size and composition (no. of males, females, adults, juveniles, sub-adults). Seasons were noted as Monsoon, Pre-Monsoon and Post-Monsoon. Arrival of Monsoon rains marks a remarkable transition of seasons on Indian subcontinent. Monsoon season is associated with greater availability of water, food and sowing of kharif crops. Monsoon season for Morachi Chincholi and Nashik is June to September while for Rajasthan it is July-September as Monsoon rains start in Rajasthan later than the other two sites. Post Monsoon season is October-January, while Pre-Monsoon season is February to May for Morachi Chincholi and Nashik, February to June for Rajasthan.

### Detection of intestinal parasites from faecal samples

Faecal samples of Indian peafowl were collected using clean forceps from food provision and non-provision areas across the three seasons (Pre-Monsoon, Monsoon and Post-Monsoon) from the field sites. Samples collected at the food provision sites were within 2m radius of the identified area where food is offered to the peafowl throughout the year. The appearance and consistency of the sample was noted down along with the GPS co-ordinates of the location where it was collected. The sample was divided roughly in two equal parts - one part in pre-weighted sample bottle containing 5ml saline and another part in pre-weighted empty bottle. The bottles were weighed again to estimate the weight of faecal sample. Faecal samples of peafowl were subjected to two methods of analysis for estimating parasitic load and diversity.

#### Method 1

Samples collected in physiological saline (0.83 %-0.9 %) were further processed with 2 % Potassium dichromate (K_2_Cr_2_O_7_) to enhance sporulation of cyst forming parasites. Then parasites were isolated using Sheather’s Sugar Flotation method (Duszynski and Wilber 1997).

#### Method 2

Sample was fixed using 10% formalin and no sporulation agent was used. Further the sample was washed with tap water, centrifuged at 2000 rpm for 5 minutes and re-suspended in 2.79M Zinc Sulfate hepta-hydrate (ZnSO_4_). The sample was again centrifuged at 2000 rpm for 5minutes (Watve and Sukumar 1992).

Samples processed in K_2_Cr_2_O_7_ enhance the sporulation of cyst forming parasites while the samples fixed in formalin might show both cyst and non-cyst forming parasites. 75-80 micro-litre of processed sample thus obtained were subjected to microscopy for quantification of parasite load and photo-documentation of parasite types using Magvision software. In the beginning, the parasites were screened in the microscopic fields ranging from 11 to 20, to see if number of fields affects the counts of parasites. As there was no correlation between number of microscopic fields sampled and parasite load (Pearson’s correlation, r=0.08, p=0.19), 20 microscopic fields were screened for each sample thereafter. Parasite load was quantified by sampling 20 fields under the microscope for each method respectively. Parasites were identified at the broader levels-group, family, class and phylum-wherever possible using references of identified parasites in literature (Snore 1939; Atkinson et al. 2008; Schoener et al. 2012; Jaiswal et al. 2013). Photos with uncertain identity were not included in the analysis.

### Statistical Analysis

All data were analysed using statistical software STATISTICA™ version 13.2 (Dell Inc. 2016). Group sizes were compared across habitats, field sites (Morachi Chincholi, Nashik, Rajasthan), time of day (morning, evening), and seasons (Pre-Monsoon, Monsoon and Post-Monsoon) using non-parametric tests as the group size data did not follow normal distribution. Parasite prevalence was calculated as (presence or absence of parasites/total no. of samples screened*100) and was not different across methods (K_2_Cr_2_O_7_=74% and ZnSO_4_=75%, N=185). Hence, for calculating parasite prevalence, presence/ absence data of parasites detected using both methods were pooled and prevalence was compared across field sites (Morachi Chincholi, Nashik and Rajasthan), season (Pre-Monsoon, Monsoon and Post-Monsoon) and provisioning status (presence/absence of feeding site). Parasite load was calculated as count of parasites in 20 microscopic fields for both the methods respectively. As parasite load was not different across methods (Wilcoxon Matched Pair tests, p=0.74, N=185), it was also pooled for both the methods and calculated by using the following formula:

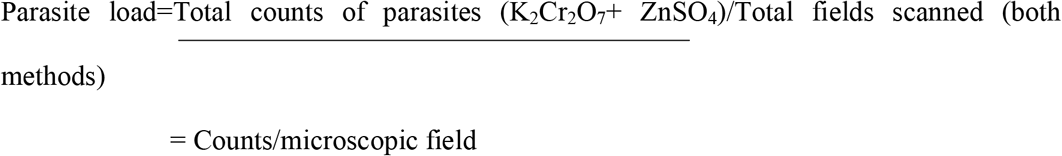

Further, the parasite load was compared across field sites, season and provisioning. Based on the presence/ absence data obtained from the photo-documentation, parasite types were identified till family level or phylum level and categorized into following groups: Eimerideae, Nematode, Cestode, Trematode and Helminth. Parasite types identified in the study can be viewed in Fig 2 and a brief summary about identification characteristics and mode of transmission is given in the Table 2.

**Table 2.** Brief summary of identification characteristics of Parasite types.

| **Parasite type** | **Identification characteristics** | **Life cycle and Mode of transmission** |
| --- | --- | --- |
| 1) Eimerideae | Typically an oocyst consisting non-sporulated or sporulated sporocysts depending on the stage of development (Atkinson et al. 2008). Size ranges around 1µm in breadth to 2µm in length when observed under the microscope with the magnification of 400X. | Oocysts shed in the faeces of infected host are taken up through food and water by non-infected individuals. The oocyst contains the infective sporozoites and these are released when the oocyst reaches the small intestine (Atkinson et al. 2008). |
| 2) Nematode | Nematode eggs in fresh faeces consist of a shell lined a semi-permeable membrane. This shell membrane may enclose one or more cells or fully grown embryo. (Snore 1939). Size ranges around 3µm in breadth and 6µm in length when observed under the microscope with the magnification of 400X. | Nematodes infect their hosts either through egg ingestion or through intermediate co-host. (Leung and Koprivnikar 2016). |
| 3) Cestode | Cestode eggs consist of two membranes. Outer membrane is rough while inner membrane is smooth and harbours a developing embryo. (Schoener et al. 2012) Size ranges around 6µm in breadth and 7µm in length when observed under the microscope with the magnification of 400X. | Cestodes follow multiple host life cycles. (Gunn and Pitt 2012). |
| 4) Trematode | The eggs are ellipsoidal in shape with a single layered membrane. Size ranges around 3µm in breadth to 5µm in length when observed under the microscope with magnification of 400X. | Trematodes follow a complex life cycle that includes one or two intermediate hosts, first of which is invariably a mollusc which can be ingested by other host (Gunn and Pitt 2012). |
| 5) Helminth | Eggs that could not be identified on the finer levels like Nematode, Cestode or Trematode were labelled as Helminth | Helminths follow multiple host life cycle and include one or two intermediate hosts. |
|  | egg. Size ranges around 2µm in breadth and 3µm in length when observed under the microscope with the magnification of 400X. | (Gunn and Pitt 2012). |

**Fig 2.**
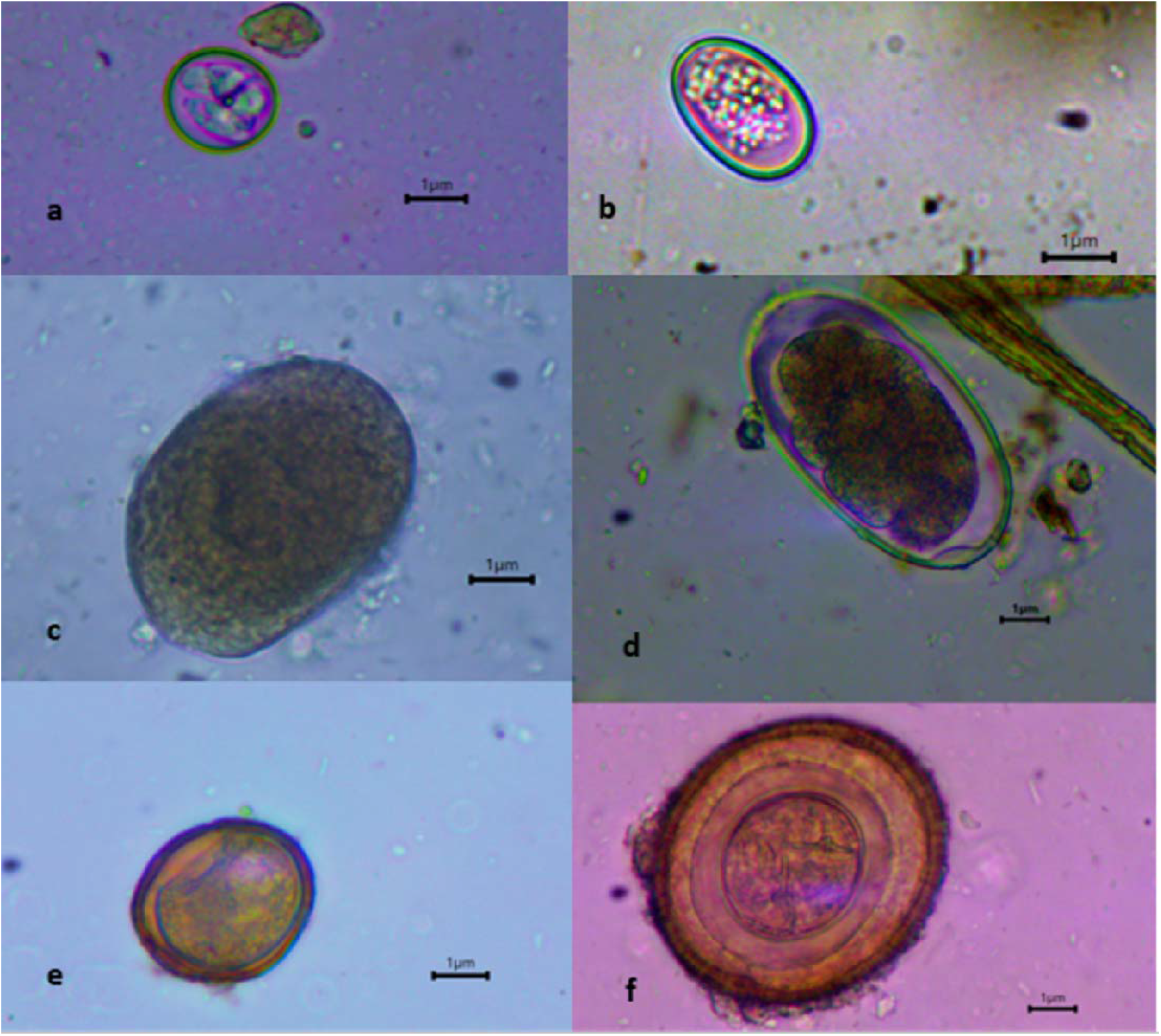
Representative photos of parasite types identified from faecal samples of Indian peafowl (a) shows sporulated Eimerideae and (b) shows non-sporulated Eimerideae, (c), (d), (e) and (f) represent Trematode, Nematode, Helminth and Cestode, respectively. Scale bars in all panels represent 1µ size

## Results

### Group size

Group size of Indian peafowl ranged from 1-14 individuals, (665 groups observed). 170 out of 665 groups observed (26%) comprised of 3 or more individuals. Ninety-nine groups contained just two individuals (15%) while 396 single individuals (59%) were observed during the study period. 57 out of 99 pairs observed (58%) had adult males as part of the pair. Out of 396 single individuals, 314 individuals were adult males (80 %), 54 females (14 %), 26 sub-adult males (6.5%) and two juveniles (0.5 %). Thus, adult males are mostly seen to live on their own or in pairs while, females, sub adult males and juveniles mostly live in groups of two or more individuals.

Group size was found to be similar across our field sites (Median test, Chi Square=1.99, df =2, p=0.36). Time of day (morning or evening) did not influence the size of group (Mann-Whitney U test, p=0.08), which indicates that the groups might not be momentary assemblages of individuals for the purpose of feeding or movement but relatively long-term associations of at least a few days/ months. The distribution of group size data across the seasons was positively skewed (Fig 3). The group size changed significantly with season (Median test, Chi square=11.44, df =2, p=0.0024). Multiple pair-wise comparisons showed that Pre-Monsoon (N=314) groups were significantly greater in size compared to groups during Monsoon season (N=238) (*p* = 0.0024), while group sizes in Post-Monsoon season (N=68) were comparable with Pre-Monsoon and Monsoon seasons (p=0.77).

**Fig 3.**
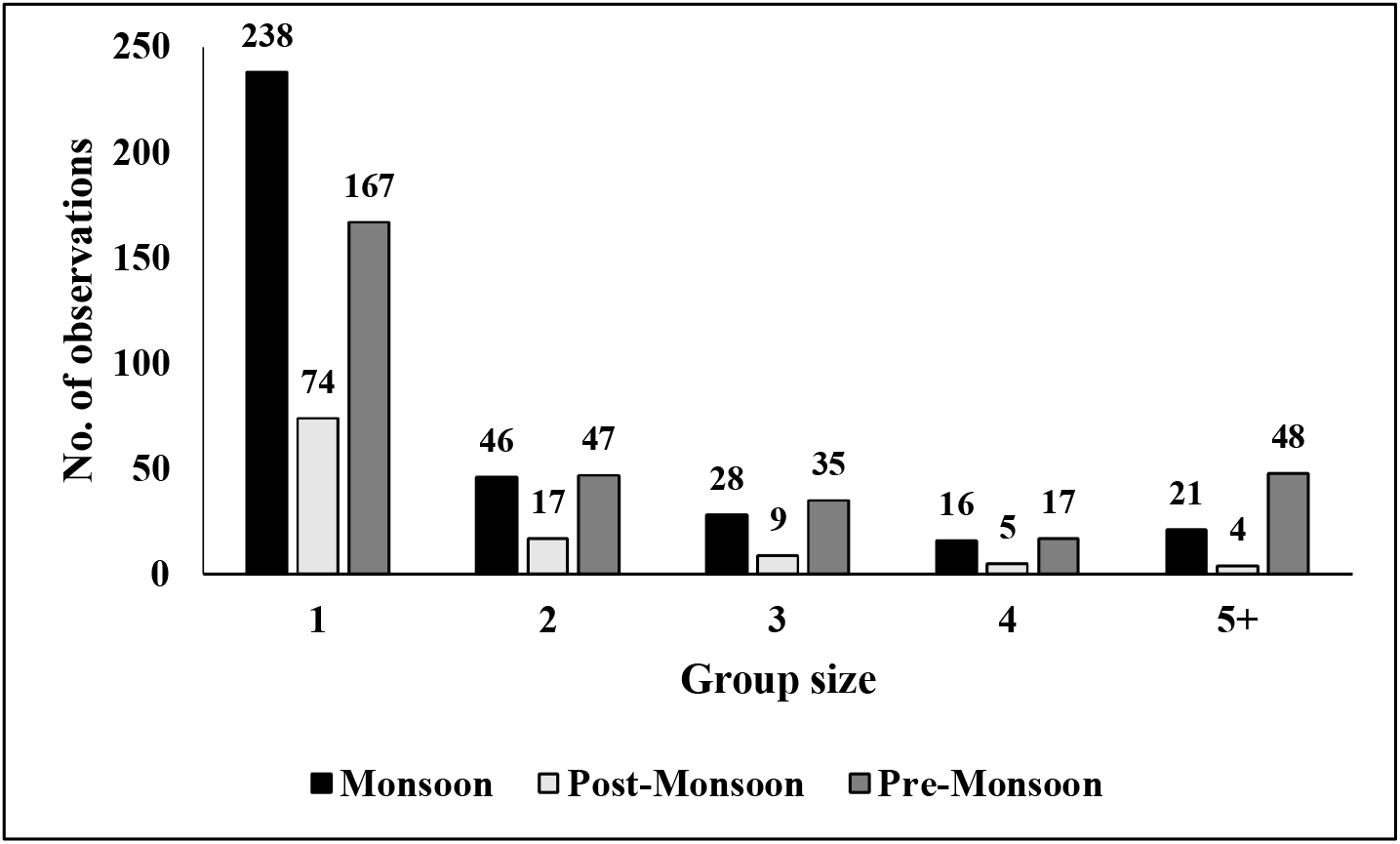
Frequency Distribution depicting Group size of Indian peafowl across Monsoon (Max. group size=8, Median= 1), Post-Monsoon (Max. group size=9, Median= 1) and Pre-Monsoon (Max. group size=14, Median=1) seasons

### Intestinal Parasites

Parasite prevalence and load (counts/microscopic field) of Indian peafowl changed according to the season. Parasite load (counts/microscopic field) ranged from 0 to 25.54 with Median=0.63 (244 samples screened). The distribution of data across the seasons (Fig 4) and provisioning (Fig 5) was positively skewed. Parasite prevalence was the least (64%) in all study populations in the Pre-Monsoon season (N=97), which increased to 91 % prevalence in Monsoon season (N=81) and 92 % in Post-Monsoon season (N=66). Similar seasonal trend was seen for parasite load, where parasite load was significantly lower in Pre-Monsoon (N=97) season compared to Monsoon (N=81) season (Median test, Chi Square=16.54, df =2, p=0.0003). Multiple comparison followed by Median test showed comparable parasite load between Post-Monsoon (N=66) and Monsoon (N=81) (p=1.0). Parasite prevalence in samples collected at food provision sites (85%, N=113) was comparable with samples collected at non-provisioned sites (78%, N=131). However, parasite load was significantly higher in faecal samples collected near the food provision sites (N=113) compared to non-provision sites (N=131) (Mann-Whitney U test, *p* = 0.004). Parasite prevalence and load was not different at field sites-Morachi Chincholi (N=116), Rajasthan (N=65) and Nashik (N=63) (Median test, Chi Square=2.556, df =2, p=0.2786).

**Fig 4.**
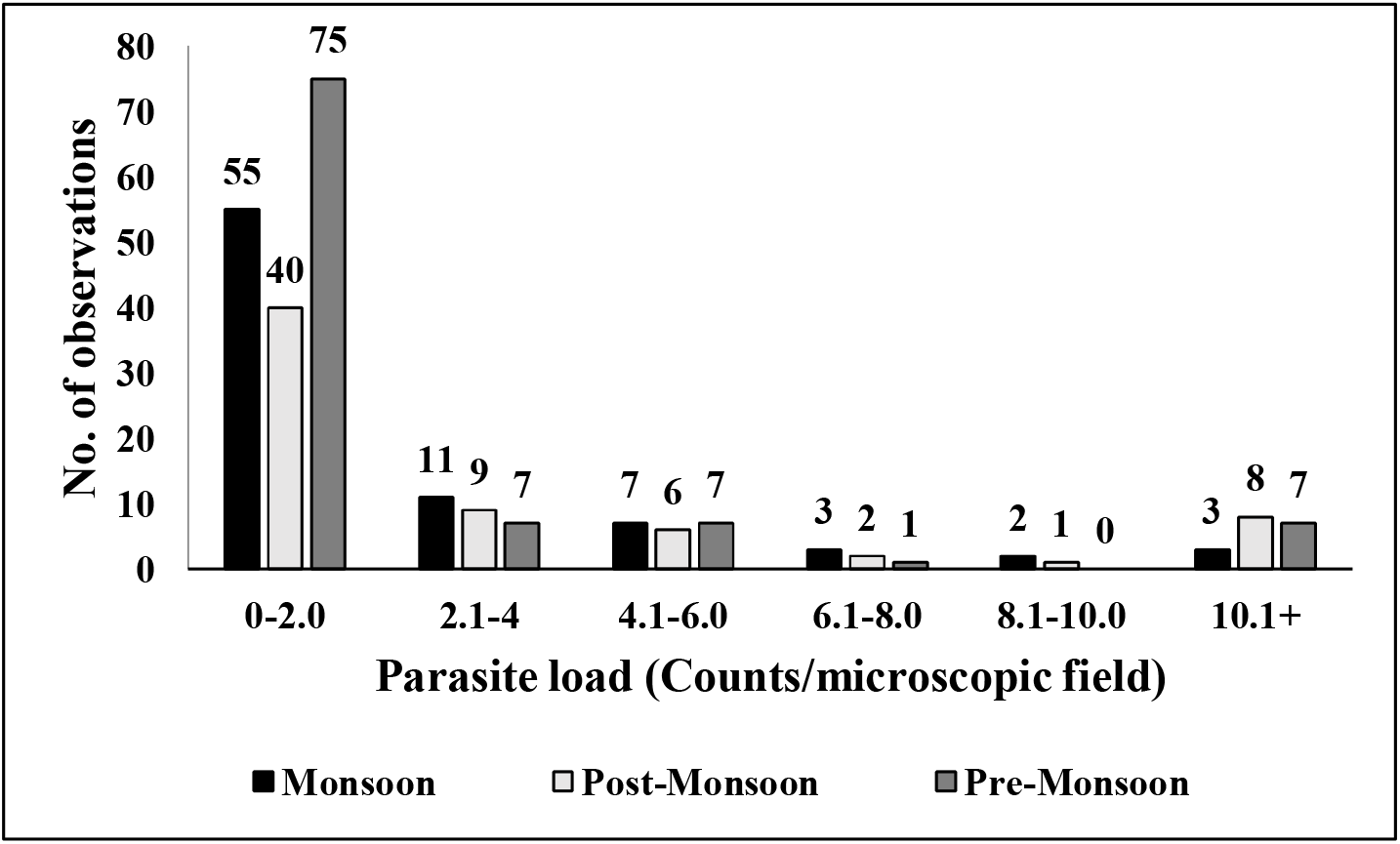
Frequency Distribution depicting Parasite load (Counts/microscopic fields) of Indian peafowl across Monsoon (Max. count= 16, Median=1.05), Post-Monsoon (Max. count= 24.62, Median=1.5) and Pre-Monsoon (Max. count= 25.54, Median=0.125) seasons

**Fig 5.**
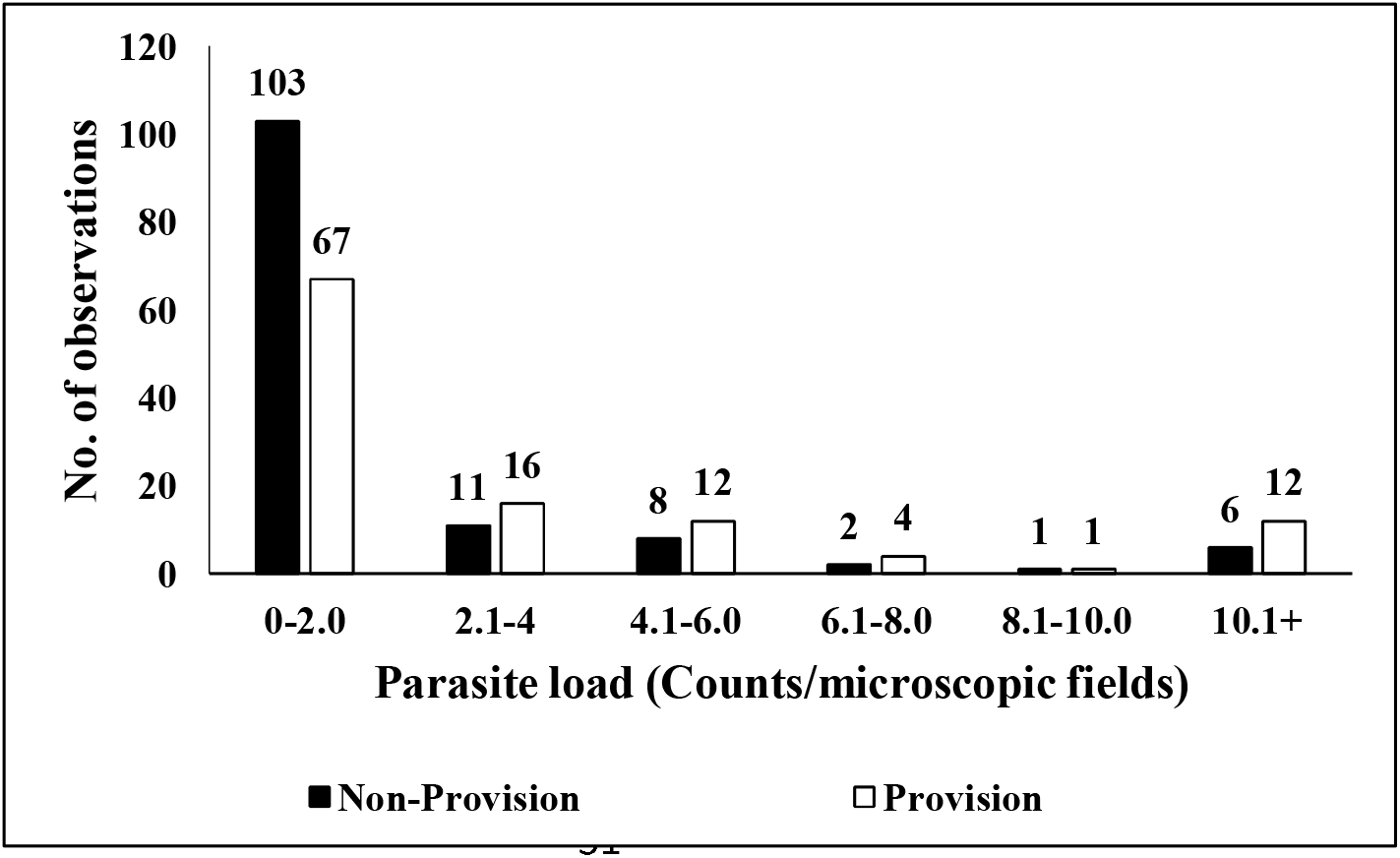
Frequency Distribution depicting Parasite load (Counts/microscopic fields) at provision (Max. count=25.54, Median=1.11) and non-provision sites (Max. count=16, Median=0.45)

The parasite types identified were grouped as Eimerideae, Nematodes, Cestodes, Trematodes and Helminths. Based on the presence / absence data collected from the samples, the percent prevalence of each parasite type was calculated as (number of positive samples for a parasite type)/ (Total no. of samples)*100. The K_2_Cr_2_O_7_ method was qualitatively better for detecting and identifying parasites belonging to family Eimerideae based on the structure of cysts compared to ZnSO_4_ method. Nematodes were detected with higher percentage in samples processed using ZNSO_4_ (20%) compared to K_2_Cr_2_O_7_ (8%).

Pre-Monsoon and Post-Monsoon seasons had lower percentage prevalence (62% and 65% respectively) for Eimerideae as compared to Monsoon (84%). (Fig 6). Cestodes were also in higher percentage in Post-Monsoon (47%) season in comparison to Monsoon (32%) and Pre-Monsoon season (20%) (Fig 6). Nematodes showed higher percentage prevalence during Post-Monsoon (35%) season as compared to Monsoon (22%) and Pre-Monsoon (20%) season (Fig 6). Cestodes were found in higher percentages at non-provision (40%) sites as compared to provision (20%) sites (Fig 7). Across the field sites, Eimerideae were detected in higher percentages in Morachi Chincholi (62%) as compared to Rajasthan (42%) and Nashik (42%) (Fig 7). Cestodes were detected in lower percentages at Rajasthan (7%) as compared to Morachi Chincholi (35 %) and Nashik (26 %) (Fig 8).

**Fig 6.**
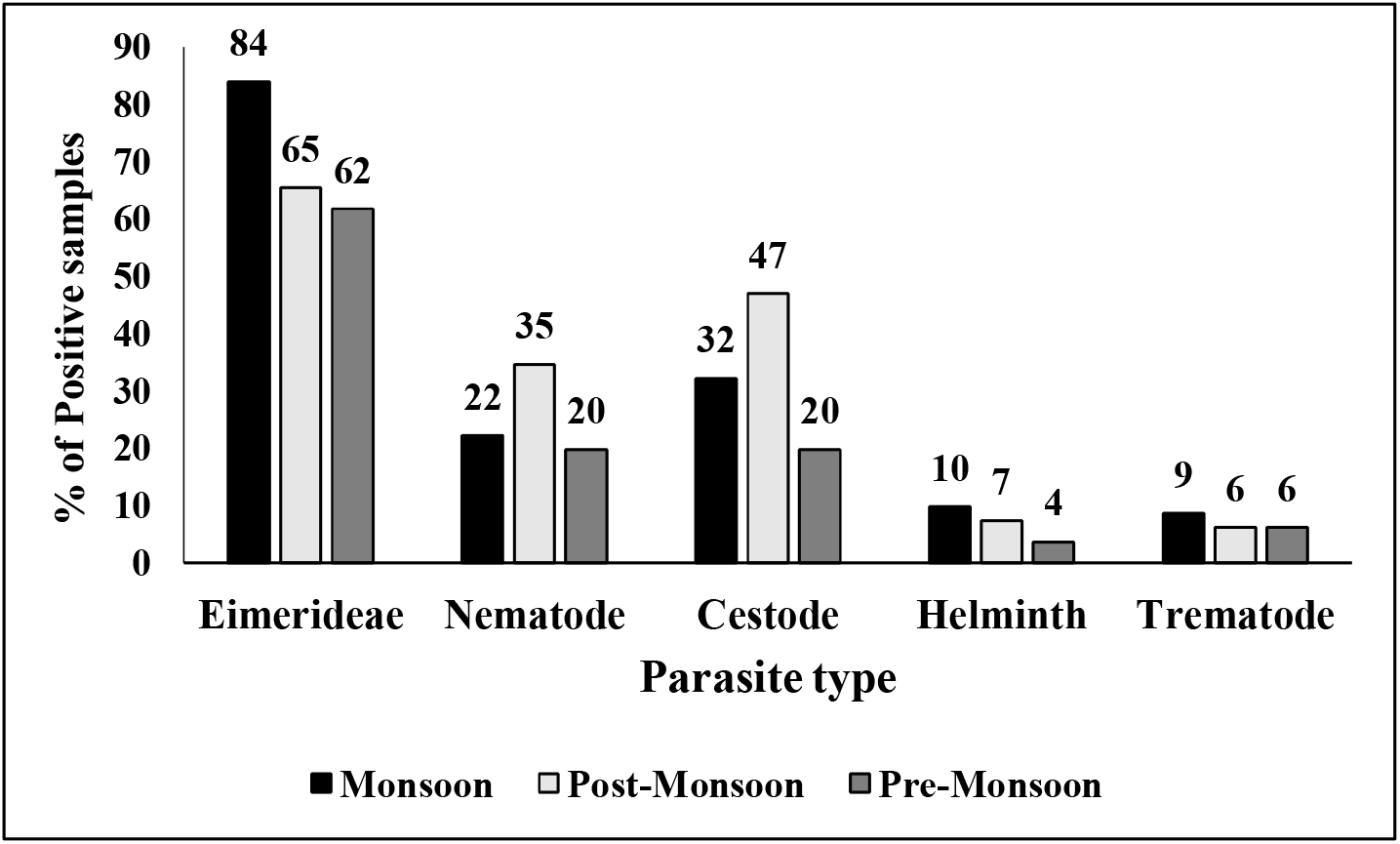
Percentage prevalence for Parasite types-Eimerideae, Nematode, Cestode, Helminth and Trematode across Monsoon, Post-Monsoon and Pre-Monsoon seasons

**Fig 7.**
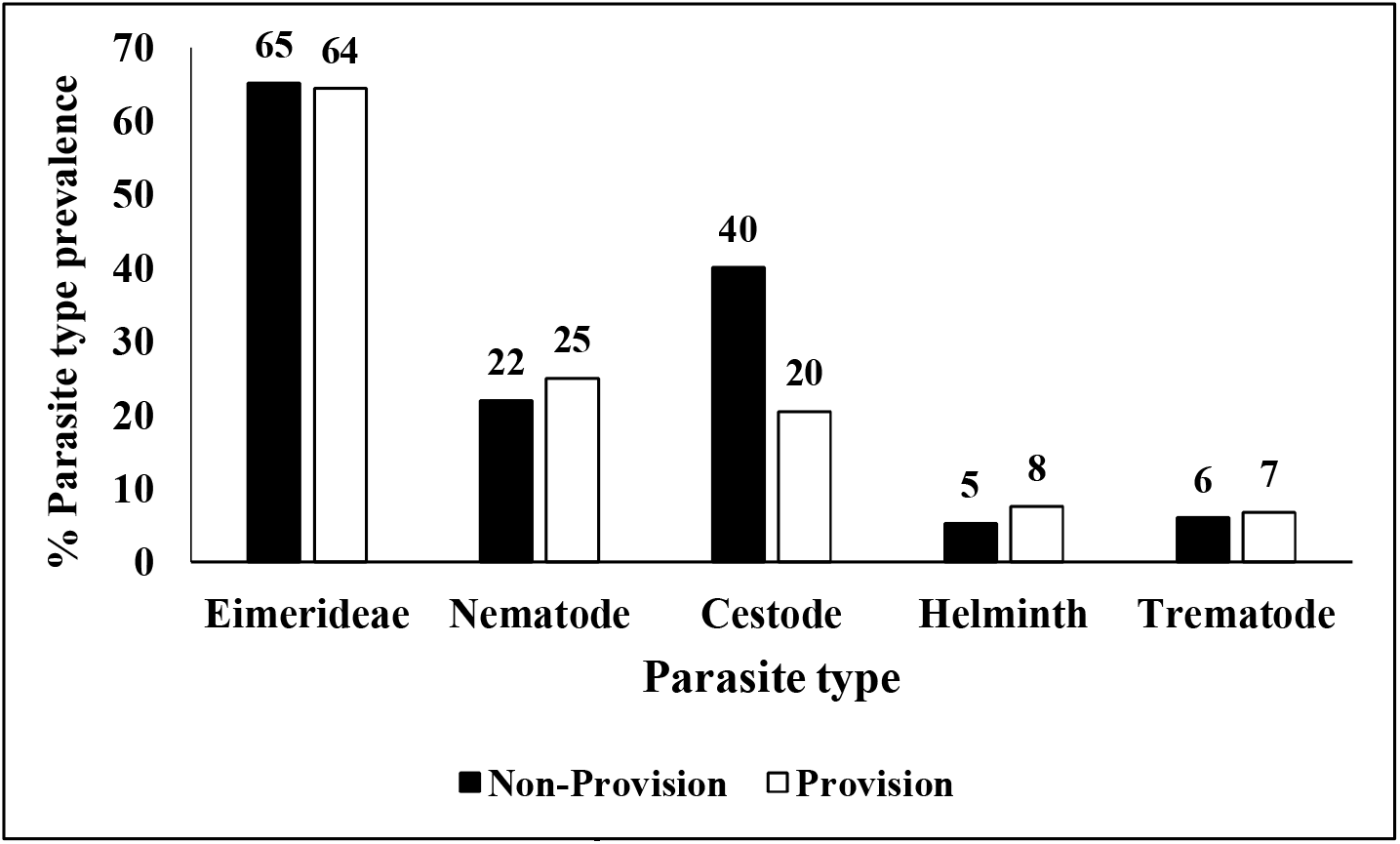
Percentage prevalence for Parasite types-Eimerideae, Nematode, Cestode, Helminth and Trematode at non-provision and provision sites

**Fig 8.**
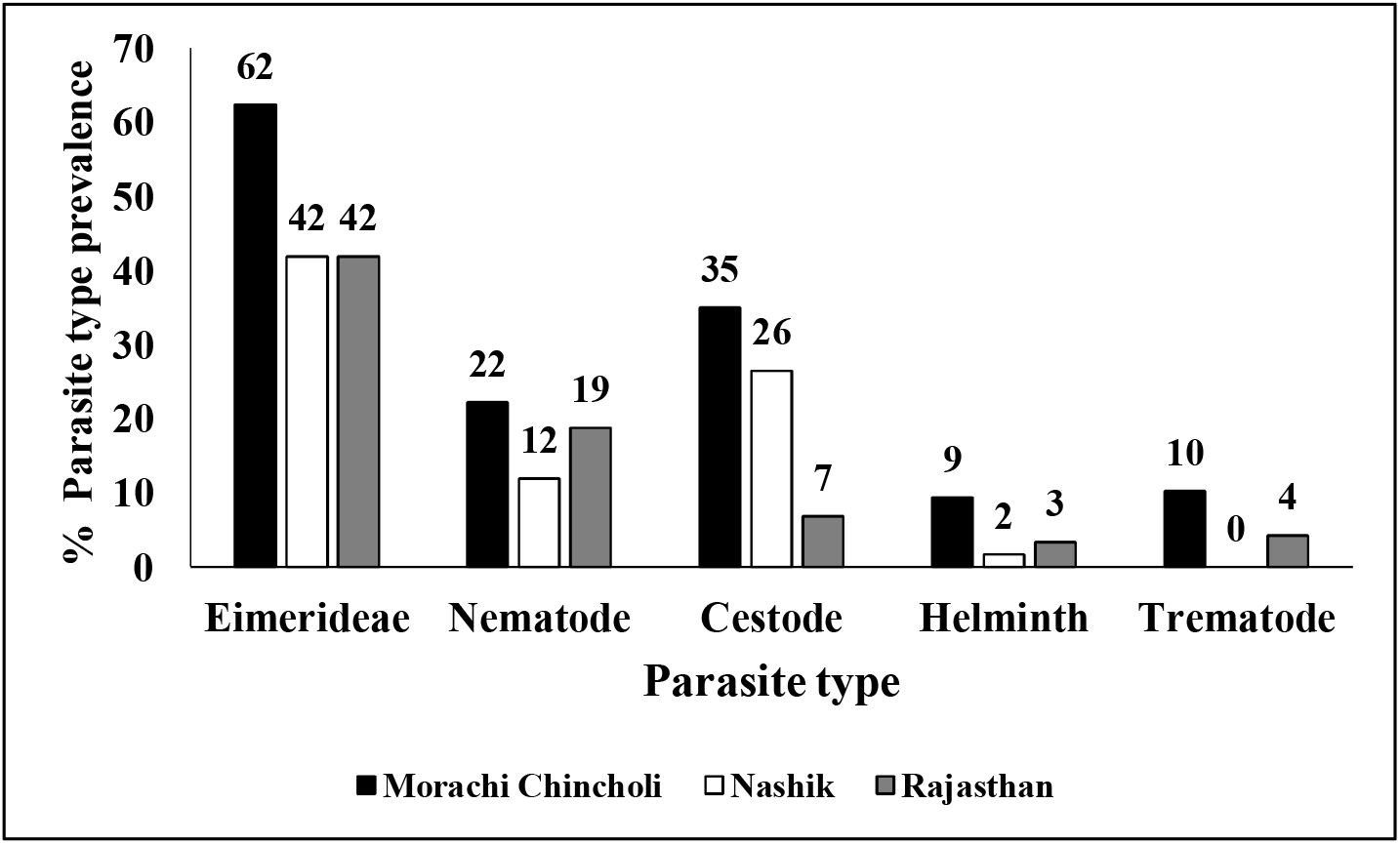
Percentage prevalence for Parasite types-Eimerideae, Nematode, Cestode, Helminth and Trematode across Morachi Chincholi, Nashik and Rajasthan

## Discussion

### Effect of season on group size and intestinal parasites of Indian peafowl

Peafowls showed larger group size in Pre-Monsoon season as compared to Monsoon and Post-Monsoon seasons. Pre-Monsoon season consists of dry hot summers at all three field sites. During the summers as vegetation dries up, peafowls can be easily spotted at large distances by their predators/intruders. In Morachi Chincholi and Nashik, feral dogs are often their predators while caracals, feral dogs and tigers are said to be the predators of Indian peafowl in Rajasthan (anecdotal). Remaining in larger groups in Pre-Monsoon season may, thus, provide safety to them from predators/ intruders or simply reduce the risk of predation due to dilution effect (Foster and Treherne 1981; Creswell 1994). In another study in Redshanks, the probability of attack by raptors was lesser in individuals staying in larger flocks as compared to individuals staying in smaller flocks (Cresswell 1994). Group size dynamics was driven by tidal cycles in Light Bellied Geese where their flock size increased with the predation risk (Inger et al. 2006). As reported in previous studies on various other study systems, there are also costs associated with living in a group such as competition for food (Krause and Ruxton, 2002) with other members of the group and higher chances of disease transmission (Hochberg 1991). Larger groups may become more conspicuous to predators (Cresswell 1994; Krause and Ruxton 2002).

In spite of these potential costs, the peafowls in our study populations were seen in larger groups during Pre-Monsoon season at all three field sites. When the Monsoon commences, the larger peafowl groups start splitting. There might be two possible reasons why peafowl groups split:

1. Newly sown crops/ sprouts, insects, worms become abundant everywhere soon after the rains start in June. So, food and water availability are not restricted to food provision sites anymore, in contrast to dry hot summer months of Pre-Monsoon season. There is still good visibility for about two months till the crops/ grasses/ vegetation grow enough. So being in a group can still offer protection against predators, however, benefits of getting varied food which is now available everywhere may override the costs of leaving the group.

2. Breeding males break off from larger groups and establish their display territories (if they haven’t already established such territory since April-May). Since they have to defend these display territories throughout the breeding season (till Sept-October, when Monsoon/ wet season also ends), breeding males can no longer afford to be part of larger groups. The peak of breeding season is June-July-August, when we mostly see solitary breeding males and groups of females + sub-adult males (See supplementary data). So the fission-fusion of groups is also related to their behavioural changes during breeding season.

Interestingly, our parasite data indicated that parasite prevalence and load in our study populations were in fact lower during Pre-Monsoon season. Thus, peafowl populations seem to be at lower risk of intestinal disease transmission in spite of being in larger groups during Pre-Monsoon season. All the parasites detected from faecal samples are intestinal parasites. Most likely their transmission happens via faecal-oral route or through intermediate hosts (Table 2 and references within). Probability of transmission of intestinal parasites is usually higher during wet season, i.e., Monsoon. Thus, increased risk of disease transmission in a larger group combined with need to establish/ defend display territories may result in many males leaving the groups during Monsoon season. Similar to our results, the prevalence of endoparasites was highest in the Monsoon season (83%) in backyard poultry in North region of India (Bhatt et al. 2014). Nematode burden was the least in dry season and highest in the rainy seasons in communal land goats of Zimbabwe (Pandey et al. 1994).

Seasonal trends are also reflected in the parasite types where Pre-Monsoon season had lower percentages of Nematode and Cestode as compared to Monsoon and Post-Monsoon season. Prevalence of Eimerideae was comparable between Post-Monsoon and Pre-Monsoon but more in Monsoon season. As Eimerideae are chiefly transmitted to next host via faecal oral route (Atkinson et al., 2008) and wet season (Monsoon) is more conducive for such transmission, Eimerideae might be more prevalent during Monsoon season compared to Pre-Monsoon and Post-Monsoon seasons.

If indeed there is lower risk of disease transmission during Pre-Monsoon season, peafowls could get benefits in terms of lower parasite transmission and reduced risk of predation when they were seen in larger groups during Pre-Monsoon season. Smaller group sizes in Monsoon and Post-Monsoon season when parasite prevalence/load is greater, might be a strategy to control parasite transmission in Indian peafowl populations. Control of parasite transmission may be important in determining the integrity of social group composition and regulation of group size (Freeland 1979). This temporal pattern of fission-fusion of groups in Indian peafowl may, thus, be a result of behavioural strategies to tackle ecological stressors such as predation and parasite transmission.

Beyond a simple dependence on group size, however, recent work in the field of network epidemiology has shown that infectious disease spread largely depends on the organization of infection-spreading interactions between individuals (Sah et al. 2017). Group composition/ organization in Indian peafowl populations consisted of adult males, who were mostly seen to live on their own or in pairs while, females, sub adult males and juveniles were seen in groups of two or more individuals. But we were not able to study the network epidemiology in those groups, as we did not have the individual identity of members of a group. Therefore, further studies might be needed to explore how group composition and social interactions within and across larger groups may influence disease transmission.

### Effect of provisioning on intestinal parasites of Indian peafowl

Samples found at food provisioning sites had significantly higher parasite load than non-provision sites. Percentage prevalence of parasite types-Eimerideae, Nematode, Helminth and Trematode was comparable at provisioned and non-provisioned sites, except for Cestode which was higher in percentage at non-provisioned sites (Fig.7.). Higher parasite load at provisioning sites may be due to aggregation of groups at provisioning sites and the increased risk of transmission of parasites. Similar trend was observed in racoon populations, where the resource distribution was altered through experimental manipulation to enhance aggregation of individuals. This resulted in significant increase of prevalence (up to 54%) of the parasitic nematode *Baylisascaris procyonis* in the experimental population (Gompper and Wright 2005). Similarly, the brucellosis infection was highly correlated with the duration and aggregation of individuals at the feeding grounds in elk populations (Cross et al. 2007). Nutritional quality of provisioned food may also affect the parasite load in the host. For example, supplemental feeding of rock iguanas by tourists in the Bahamas with carbohydrate-rich foods such as cereals and grapes was associated with altered nutritional status and increased hookworm burdens (Knapp et al. 2013). On the contrary, in lace monitor, supplementary feeding in form of accidental urban waste, improved nutrition and lowered intensity of pathogen *Haemogregarina varanicola* (Jessop et al. 2012).

In this study, individuals were not marked and it is very difficult to identify and keep track of individuals by sight. Therefore, there is a possibility that faecal samples of some individuals may have been sampled repeatedly at provision and non-provision sites and across seasons. However, there was no bias in sampling with respect to healthy or infected individuals as the data was collected across 3 years and all areas at each field site were sampled consistently across all seasons. Apart from food provisioning and season, other factors such as stage of infection within the host, immunity and stress levels of the host may influence the number of eggs or cysts released into the faecal samples. However, the estimated parasite load can still give us valuable information about how many individuals in the population are capable of spreading the infection and the health status of the infected individuals at a given time. This information will be further useful to address how parasite load and the spread of infection can influence the group size dynamics.

## Conclusions

In many group-living species, groups may merge or split as they move in their environment. In this study, Indian peafowl (*Pavo cristatus*) exhibited fission-fusion dynamics in their group sizes across seasons. These temporal changes in group size were associated with the dynamics of intestinal parasite prevalence and load in those seasons. Aggregation of groups of peafowl at food provision sites matched with higher parasite prevalence and load. Most of the earlier studies on intestinal parasites in Indian peafowl have focused on classifying and describing various types of parasites. Our study goes one step further in exploring association of intestinal parasites in Indian peafowls with group size dynamics, seasonal changes as well as anthropogenic factors such as food provisioning. Thus, the study furthers our understanding of factors that shape the grouping behaviour of a mostly ground dwelling bird species in the context of ecological stressors such as parasite transmission and predation.

## Supporting information

Group size data

## Acknowledgements

Thanks to Maharashtra Education Society’s Abasaheb Garware College, Pune for hosting the Ramalingaswami Re-entry Fellowship to DAP and providing the infrastructure and administrative support. We are thankful to Late Mr. Biswaroop Raha for suggesting field sites in Nashik, Dr. Dharmendra Khandal and volunteers of Tiger Watch who extended support for field work in Rajasthan. Authors also appreciate the support and hospitality of Morachi Chincholi residents during field work. We are also thankful to Rupesh Gawade, Prasad Gond, Nishant Zazam, Vishal Varma, Apeksha Dharshetkar, Priyanka Bansode, Eshan Pahade, Ankita Divekar and Akash Dubey and Vedanti Mahimkar for helping with data collection on field. We want to thank the anonymous reviewers for their valuable suggestions which increased the quality of this manuscript.

## Declarations

### Ethics approval

No live animals/ samples were handled during this study. The study was conducted outside protected areas. Therefore, the work did not require special permissions or approval.

### Consent to participate

**Not applicable**

### Consent for publication

**Not applicable**

### Availability of data and material

The data is made available as electronic supplementary material with this manuscript.

### Authors contributions

DAP conceived the study, designed methodology, collected data and helped in manuscript preparation. PD designed the methodology, collected data, analysed the data and prepared the manuscript. PM collected and analysed the data.

## References

Atkinson CT, Thomas NJ, Hunter DB (2008) Parasitic Diseases of Wild Birds. Wiley-Blackwell, U.S.A

Aureli F, Schaffner CM, Boesch C, Bearder SK, Call J, Chapman CA, Connor R, Fiore A, Dunbar RIM (2008) Fission-Fusion Dynamics. Curr Anthropol 49:627–654. doi: 10.1086/586708

Bhatt S, Khajuria JK, Katoch R, Dhama K (2014) Prevalence of endoparasites in Backyard poultry in North Indian Region: A Performance Based Assessment Study. Asian J Anim Vet Adv 9:479–488. doi: 10.3923/ajava.2014.479.488

Beauchamp G (2010) Relationship between distance to cover, vigilance and group size in staging flocks of Semipalmated sandpipers. J Ethol 116:645–652.doi: 10.1111/j.1439-0310.2010.01778.x

Burton M, Burton R (2002) International Wildlife Encyclopedia. Marshall Cavendish, New York.

Caillaud D, Levrero F, Cristescu R, Gatti S, Dewas M, Douadi M, Gautier-Hion A, Raymond M, Meynard N (2006) Gorilla susceptibility to Ebola virus: The cost of sociality. Curr Biol Magazine R 489. doi :10.1016/j.cub.2006.06.017

Clark CW, Mangel M (1986) The Evolutionary Advantages of Group Foraging. Theor Popul Biol 30: 45–47. doi: 10.1016/0040-5809(86)90024-9

Clutton-Brock TH, Parker GA (1995) Sexual coercion in animal societies. Anim Behav 49: 1345–1365. doi: 10.1006/anbe.1995.0166.

Couzin ID (2006) Behavioral ecology: social organization in fission–fusion societies. Curr Biol R 169–R171. doi:10.1016/j.cub.2006.02.042

Cresswell W (1994) Flocking is an effective anti-predation strategy in redshanks, Tringa totanus. Anim Behav 47: 433–442.doi: 10.1006/anbe.1994.1057

Croft DP, Krause J, James R (2004) Social networks in the guppy (Poecilia reticulata). Proc R Soc B 271: 516–519.doi: 10.1098/rsbl.2004.0206

Cross PC, Edwards WH, Scurlock BM, Maichak EJ, Rogerson JD (2007) Effects of management and climate on elk brucellosis in the Greater Yellowstone Ecosystem. Ecol Appl 17:957–964. doi:10.1890/06-1603

Dusyzanski D, Wilber PG (1997) A Guideline for the Preparation of Species Descriptions in the Eimeriidae. J Parasitol 83: 333–336. doi: 10.2307/3284470

FishlockV, Lee PC (2013) Forest elephants: fission–fusion and social arenas. Anim Behav 85: 357–363. doi: 10.1016/j.anbehav.2012.11.004

Fleischer RC (1983) Relationships between tidal oscillations and Ruddy Turnstone flocking, foraging and vigilance behaviour. Condor 85: 22–29. doi: 10.2307/1367881

Foster WA, Treherne JE (1981) Evidence for the dilution effect in the selfish herd from fish predation on a marine insect. Nature 293: 466–467.doi:10.1038/293466a0

Freeland WJ (1979) Primate social groups as biological islands. Ecology 60: 719–728. doi:10.2307/1936609

Gompper ME, Wright, AN (2005) Altered prevalence of raccoon roundworm (Baylisascaris procyonis) owing to manipulated contact rates of hosts. J Zool 266: 215–219. doi:10.1017/S0952836905006813

Gunn A, Pitt SJ (2012) Parasitology An Integrated Approach. Wiley-Blackwell, UK

Hochberg M (1991) Viruses as costs to gregarious feeding behaviour in the Lepidoptera. Oikos 61: 291–296. doi: 10.2307/3545236

Inger R, Bearhop S, Robinson JA, Ruxton G (2006) Prey choice affects the trade-off balance between predation and starvation in an avian herbivore. Anim Behav 71:1335–1341. doi: 10.1016/j.anbehav.2005.08.015

Jaiswal AK, Sudan V, Shanker, D, Kumar P (2013) Endoparasitic infections in Indian peacocks (Pavo cristatus) of Veterinary College Campus, Mathura. J Parasit Disease 37: 26– 28.doi: 10.1007/s12639-012-0124-1

Jessop TS, Smissen P, Scheelings F, Dempster T (2012) Demographic and Phenotypic Effects of Human Mediated Trophic Subsidy on a Large Australian Lizard (Varanus varius): Meal Ticket or Last Supper? PLoS One 7:1–8. doi:10.1371/journal.pone.0034069

Kashima K, Ohtsuki H, Satake A (2013) Fission-fusion bat behaviour as a strategy for balancing the conflicting needs of maximizing information accuracy and minimizing infection risk. J Theor Biol 318:101–109. doi: 10.1016/j.jtbi.2012.10.034

Knapp CR, Hines KN, Zachariah TT, Perez-Heydrich C, Iverson JB, Buckner SD, Halach SC, Lattin CR, Romero ML (2013) Physiological effects of tourism and associated food provisioning in an endangered iguana. Conserv Physiol 1:1–12. doi: 10.1093/conphys/cot032

Krause J, Ruxton GD (2002) Living in Groups. New York: Oxford University Press.

Lehmann J, Boesch C (2004) To fission or to fusion: effects of community size on wild chimpanzee (Pan troglodytes verus) social organisation. Behav Ecol Sociobiol 56: 207–216. doi: 10.1007/s00265-004-0781

Leung TLF, Koprivnikar J (2016) Nematode parasite diversity in birds: the role of host ecology, life history and migration. J Anim Ecol 85: 1471–1480. doi: 10.1111/1365-2656.12581

Lutz HL,Hochachka WM, Engel JI, Bell JA, Tkach VV, Bates JM, Hackett JS, Weckstein JD (2015) Parasite prevalence corresponds to host life history in a diverse assemblage of Afrotropical birds and Haemosporidian parasites. PLoS ONE 10: 1–24. doi:10.1371/journal.pone.0121254

Nandini S, Keerthipriya P, Vidya TNC (2017) Seasonal variation in female Asian elephant social structure in Nagarhole-Bandipur, southern India. Anim Behav 134: 135–145. doi: 10.1016/j.anbehav.2017.10.012

Pandey VS, Ndao M, Kumar V (1994) Seasonal prevalence of gastrointestinal nematodes in communal land goats of from the highveld of Zimbabve. Vet Parasitol 51: 241–248. doi: 10.1016/0304-4017(94)90161-9

Paranjpe DA, Dange PM (2020) A tale of two species: human and peafowl interactions in human dominated landscape influence each other’s behaviour: Wild life near human habitation. Curr Sci 119: 670–679. 10.18520/cs/v119/i4/670-679

Popa-Lisseanu AG, Bontadina F, Mora O, IbÁñez C (2008) Highly structured fission–fusion societies in an aerial-hawking, carnivorous bat. Anim Behav 75: 471–482. doi: 10.1016/j.anbehav.2007.05.011

Powell G (1974) Experimental analysis of the social value of flocking by starlings (Sturnus vulgaris) in relation to predation and foraging. Anim Behav 22: 501–505. doi: 10.1016/S0003-3472(74)80049-7

Pulliam HR (1973) On the advantages of Flocking. J Theor Biol 38: 419–422. doi: 10.1016/0022-5193(73)90184-7

Rand MRW, Ridley MW, Lelliot AD (1984) The social organization of feral Indian peafowl. Anim Behav 32: 830–835. doi: 10.1016/S0003-3472(84)80159-1

Rimbach R, Bisanzio D, Galvis N, Link A, Di Fore A, Gillespie TR (2015) Brown spider monkeys (Ateles hybridus): a model for differentiating the role of social networks and physical contact on parasite transmission dynamics. Phil Trans R Soc B 370: 1–10. doi: 10.1098/rstb.2014.0110

Robb GN, McDonald RA, Chamberlain DE, Bearhop S (2008) Food for thought: supplementary feeding as a driver of ecological change in avian populations. Front Ecol Environ 6: 476–484. doi: 10.1890/060152

Sah P, Mann J, Bansal S (2017) Disease implications of animal social network structure: A synthesis across social systems. J Anim Ecol 87: 546–558. doi: 10.1111/1365-2656

Schoener ER, Howe L, Castro I, Alley MR (2012) Helminths in endemic, native and introduced passerines in New Zealand. N Z J Zool 39: 245–256. doi: 10.1080/03014223.2012.676992

Silk JB (2007) The adaptive value of sociality in mammalian groups. Phil Trans Biol Sci 362: 539–559. doi: 10.1098/rstb.2006.1994

Silk MJ, Croft DP, Trgenza T, Bearhop S (2014) The importance of fission–fusion social group dynamics in birds. Ibis 156: 701–715. doi: 10.1111/ibi.12191

Smith H, Frère C, Kobryn H, Bejder L (2016) Dolphin sociality, distribution and calving as important behavioural patterns informing management. Anim Conserv 19: 462–471. doi:10.1111/acv.12263

Snore DA (1939) Differentiation of eggs of various genera of Nematodes parasitic in domestic ruminants in the United States. USDA, Washington, D.C. U.S.A

Sutton N (2019) Effects of food limitation on social grouping and foraging in a fission-fusion species. Dissertation, University of Cantebury.vanSchaik CP (1999) The socioecology of fission-fusion sociality in Orangutans. Primates 40: 69–86. doi: 10.1007/BF02557703

Watve MG, Sukumar R (1992) Parasite abundance and diversity in mammals: Correlates with host ecology. Proc Natl Acad Sci U.S.A. 92: 8945–8949. doi: 10.1073/pnas.92.19.8945

Wittemeyer G, Douglas-Hamilton I, Getz WM (2005) The socioecology of elephants: analysis of the processes creating multitiered social structures. Anim Behav 69: 1357–1371. doi: 10.1016/j.anbehav.2004.08.018

