## Supplementary material for "To group or not to group: group size dynamics and intestinal parasites in Indian peafowl populations": Group size data

Supplementary Data

(B=Breeding season, NB=Non-breeding season)

(M = Monsoon, PostM = Post monsoon, PreM = Pre-monsoon)

| Sr.No. | Sample No. | Field site | B/NB | Season | Males | Sub-adult Male | Females | Juvenile | Unidentified | Total group size |
|---|---|---|---|---|---|---|---|---|---|---|
| 1 | 76 | Morachi Chincholi | B | M | 1 | | | | | 1 |
| 2 | 83 | Morachi Chincholi | B | M | | | 1 | | | 1 |
| 3 | 86 | Morachi Chincholi | B | M | 1 | | | | | 1 |
| 4 | 87 | Morachi Chincholi | B | M | 1 | | | | | 1 |
| 5 | 92 | Morachi Chincholi | B | M | 1 | | | | | 1 |
| 6 | 93 | Morachi Chincholi | B | M | 1 | | | | | 1 |
| 7 | 94 | Morachi Chincholi | B | M | 1 | | | | | 1 |
| 8 | 95 | Morachi Chincholi | B | M | 1 | | | | | 1 |
| 9 | 97 | Morachi Chincholi | B | M | 1 | | | | | 1 |
| 10 | 98 | Morachi Chincholi | B | M | 1 | | | | | 1 |
| 11 | 102 | Morachi Chincholi | B | M | | 1 | | | | 1 |
| 12 | 103 | Morachi Chincholi | B | M | 1 | | | | | 1 |
| 13 | 108 | Morachi Chincholi | B | M | 1 | | | | | 1 |
| 14 | 124 | Morachi Chincholi | B | M | 1 | | | | | 1 |
| 15 | 126 | Morachi Chincholi | B | M | 1 | | | | | 1 |
| 16 | 127 | Morachi Chincholi | B | M | 1 | | | | | 1 |
| 17 | 129 | Morachi Chincholi | B | M | 1 | | | | | 1 |
| 18 | 288 | Morachi Chincholi | B | M | 1 | | | | | 1 |
| 19 | 289 | Morachi Chincholi | B | M | 1 | | | | | 1 |
| 20 | 290 | Morachi Chincholi | B | M | 1 | | | | | 1 |
| 21 | 291 | Morachi Chincholi | B | M | 1 | | | | | 1 |
| 22 | 295 | Morachi Chincholi | B | M | | 1 | | | | 1 |
| 23 | 296 | Morachi Chincholi | B | M | 1 | | | | | 1 |
| 24 | 299 | Morachi Chincholi | B | M | 1 | | | | | 1 |
| 25 | 301 | Morachi Chincholi | B | M | 1 | | | | | 1 |

| | | | | | | | | | | |
|---|---|---|---|---|---|---|---|---|---|---|
| 26 | 302 | Morachi Chincholi | B | M | 1 | | | | | 1 |
| 27 | 308 | Morachi Chincholi | B | M | 1 | | | | | 1 |
| 28 | 311 | Morachi Chincholi | B | M | 1 | | | | | 1 |
| 29 | 315 | Morachi Chincholi | B | M | | | 1 | | | 1 |
| 30 | 316 | Morachi Chincholi | B | M | 1 | | | | | 1 |
| 31 | 317 | Morachi Chincholi | B | M | 1 | | | | | 1 |
| 32 | 318 | Morachi Chincholi | B | M | | | 1 | | | 1 |
| 33 | 320 | Morachi Chincholi | B | M | 1 | | | | | 1 |
| 34 | 321 | Morachi Chincholi | B | M | 1 | | | | | 1 |
| 35 | 322 | Morachi Chincholi | B | M | | | 1 | | | 1 |
| 36 | 324 | Morachi Chincholi | B | M | 1 | | | | | 1 |
| 37 | 330 | Morachi Chincholi | B | M | 1 | | | | | 1 |
| 38 | 331 | Morachi Chincholi | B | M | 1 | | | | | 1 |
| 39 | 332 | Morachi Chincholi | B | M | 1 | | | | | 1 |
| 40 | 333 | Morachi Chincholi | B | M | 1 | | | | | 1 |
| 41 | 334 | Morachi Chincholi | B | M | 1 | | | | | 1 |
| 42 | 335 | Morachi Chincholi | B | M | 1 | | | | | 1 |
| 43 | 337 | Morachi Chincholi | B | M | 1 | | | | | 1 |
| 44 | 339 | Morachi Chincholi | B | M | 1 | | | | | 1 |
| 45 | 344 | Morachi Chincholi | B | M | 1 | | | | | 1 |
| 46 | 345 | Morachi Chincholi | B | M | 1 | | | | | 1 |
| 47 | 346 | Morachi Chincholi | B | M | 1 | | | | | 1 |
| 48 | 347 | Morachi Chincholi | B | M | 1 | | | | | 1 |
| 49 | 348 | Morachi Chincholi | B | M | 1 | | | | | 1 |
| 50 | 349 | Morachi Chincholi | B | M | 1 | | | | | 1 |
| 51 | 353 | Morachi Chincholi | B | M | 1 | | | | | 1 |
| 52 | 354 | Morachi Chincholi | B | M | 1 | | | | | 1 |

| | | | | | | | | | | |
|---|---|---|---|---|---|---|---|---|---|---|
| 53 | 361 | Morachi Chincholi | B | M | | | 1 | | | 1 |
| 54 | 365 | Morachi Chincholi | B | M | 1 | | | | | 1 |
| 55 | 366 | Morachi Chincholi | B | M | 1 | | | | | 1 |
| 56 | 367 | Morachi Chincholi | B | M | 1 | | | | | 1 |
| 57 | 369 | Morachi Chincholi | B | M | 1 | | | | | 1 |
| 58 | 370 | Morachi Chincholi | B | M | | 1 | | | | 1 |
| 59 | 373 | Morachi Chincholi | B | M | | | 1 | | | 1 |
| 60 | 374 | Morachi Chincholi | B | M | | | 1 | | | 1 |
| 61 | 375 | Morachi Chincholi | B | M | | 1 | | | | 1 |
| 62 | 377 | Morachi Chincholi | B | M | 1 | | | | | 1 |
| 63 | 380 | Morachi Chincholi | B | M | 1 | | | | | 1 |
| 64 | 381 | Morachi Chincholi | B | M | 1 | | | | | 1 |
| 65 | 527 | Morachi Chincholi | B | M | 1 | | | | | 1 |
| 66 | 528 | Morachi Chincholi | B | M | 1 | | | | | 1 |
| 67 | 529 | Morachi Chincholi | B | M | 1 | | | | | 1 |
| 68 | 531 | Morachi Chincholi | B | M | 1 | | | | | 1 |
| 69 | 537 | Morachi Chincholi | B | M | 1 | | | | | 1 |
| 70 | 538 | Morachi Chincholi | B | M | 1 | | | | | 1 |
| 71 | 539 | Morachi Chincholi | B | M | 1 | | | | | 1 |
| 72 | 633 | Morachi Chincholi | B | M | 1 | | | | | 1 |
| 73 | 643 | Morachi Chincholi | B | M | 1 | | | | | 1 |
| 74 | 646 | Morachi Chincholi | B | M | 1 | | | | | 1 |
| 75 | 647 | Morachi Chincholi | B | M | 1 | | | | | 1 |
| 76 | 648 | Morachi Chincholi | B | M | | 1 | | | | 1 |
| 77 | 649 | Morachi Chincholi | B | M | 1 | | | | | 1 |
| 78 | 650 | Morachi Chincholi | B | M | | | 1` | | | 1 |
| 79 | 652 | Morachi Chincholi | B | M | 1 | | | | | 1 |

| | | | | | | | | | | |
|---|---|---|---|---|---|---|---|---|---|---|
| 80 | 653 | Morachi Chincholi | B | M | 1 | | | | | 1 |
| 81 | 654 | Morachi Chincholi | B | M | | 1 | | | | 1 |
| 82 | 655 | Morachi Chincholi | B | M | | | 1 | | | 1 |
| 83 | 658 | Morachi Chincholi | B | M | | | 1 | | | 1 |
| 84 | 659 | Morachi Chincholi | B | M | 1 | | | | | 1 |
| 85 | 660 | Morachi Chincholi | B | M | 1 | | | | | 1 |
| 86 | 663 | Morachi Chincholi | B | M | 1 | | | | | 1 |
| 87 | 664 | Morachi Chincholi | B | M | | | 1 | | | 1 |
| 88 | 668 | Morachi Chincholi | B | M | | 1 | | | | 1 |
| 89 | 669 | Morachi Chincholi | B | M | 1 | | | | | 1 |
| 90 | 675 | Morachi Chincholi | B | M | | | 1 | | | 1 |
| 91 | 691 | Morachi Chincholi | B | M | 1 | | | | | 1 |
| 92 | 692 | Morachi Chincholi | B | M | 1 | | | | | 1 |
| 93 | 693 | Morachi Chincholi | B | M | 1 | | | | | 1 |
| 94 | 90 | Morachi Chincholi | B | M | 1 | | | | | 1 |
| 95 | 91 | Morachi Chincholi | B | M | | | 1 | | | 1 |
| 96 | 258 | Morachi Chincholi | B | M | | | 1 | | | 1 |
| 97 | 259 | Morachi Chincholi | B | M | 1 | | | | | 1 |
| 98 | 260 | Morachi Chincholi | B | M | 1 | | | | | 1 |
| 99 | 306 | Morachi Chincholi | B | M | 1 | | | | | 1 |
| 100 | 307 | Morachi Chincholi | B | M | | 1 | | | | 1 |
| 101 | 287 | Morachi Chincholi | B | M | | | 1 | | | 1 |
| 102 | 78 | Morachi Chincholi | B | M | 1 | | | | | 1 |
| 103 | 79 | Morachi Chincholi | B | M | 1 | | | | | 1 |
| 104 | 80 | Morachi Chincholi | B | M | | 1 | | | | 1 |
| 105 | 327 | Morachi Chincholi | B | M | | 1 | | | | 1 |
| 106 | 371 | Morachi Chincholi | B | M | 1 | | | | | 1 |

| | | | | | | | | | | |
|---|---|---|---|---|---|---|---|---|---|---|
| 107 | 372 | Morachi Chincholi | B | M | 1 | | | | | 1 |
| 108 | 625 | Morachi Chincholi | B | M | 1 | | | | | 1 |
| 109 | 626 | Morachi Chincholi | B | M | | | 1 | | | 1 |
| 110 | 627 | Morachi Chincholi | B | M | 1 | | | | | 1 |
| 111 | 628 | Morachi Chincholi | B | M | 1 | | | | | 1 |
| 112 | 629 | Morachi Chincholi | B | M | 1 | | | | | 1 |
| 113 | 631 | Morachi Chincholi | B | M | 1 | | | | | 1 |
| 114 | 635 | Morachi Chincholi | B | M | 1 | | | | | 1 |
| 115 | 636 | Morachi Chincholi | B | M | 1 | | | | | 1 |
| 116 | 638 | Morachi Chincholi | B | M | 1 | | | | | 1 |
| 117 | 639 | Morachi Chincholi | B | M | | | 1 | | | 1 |
| 118 | 640 | Morachi Chincholi | B | M | 1 | | | | | 1 |
| 119 | 641 | Morachi Chincholi | B | M | 1 | | | | | 1 |
| 120 | 642 | Morachi Chincholi | B | M | 1 | | | | | 1 |
| 121 | 77 | Morachi Chincholi | B | M | 1 | | | | | 1 |
| 122 | 81 | Morachi Chincholi | B | M | 1 | | | | | 1 |
| 123 | 99 | Morachi Chincholi | B | M | 1 | | | | | 1 |
| 124 | 356 | Morachi Chincholi | B | M | | | 1 | | | 1 |
| 125 | 357 | Morachi Chincholi | B | M | | | 1 | | | 1 |
| 126 | 624 | Morachi Chincholi | B | M | 1 | | | | | 1 |
| 127 | 667 | Morachi Chincholi | B | M | 1 | | | | | 1 |
| 128 | 670 | Morachi Chincholi | B | M | 1 | | | | | 1 |
| 129 | 85 | Morachi Chincholi | B | M | 1 | | | | | 1 |
| 130 | 309 | Morachi Chincholi | B | M | | 1 | | | | 1 |
| 131 | 312 | Morachi Chincholi | B | M | 1 | | | | | 1 |
| 132 | 360 | Morachi Chincholi | B | M | | | 1 | | | 1 |
| 133 | 362 | Morachi Chincholi | B | M | 1 | | | | | 1 |

| | | | | | | | | | | |
|---|---|---|---|---|---|---|---|---|---|---|
| 134 | 634 | Morachi Chincholi | B | M | 1 | | | | | 1 |
| 135 | 96 | Morachi Chincholi | B | M | 1 | | | | | 1 |
| 136 | 104 | Morachi Chincholi | B | M | | | 1 | | | 1 |
| 137 | 340 | Morachi Chincholi | B | M | 1 | | | | | 1 |
| 138 | 341 | Morachi Chincholi | B | M | 1 | | | | | 1 |
| 139 | 342 | Morachi Chincholi | B | M | 1 | | | | | 1 |
| 140 | 363 | Morachi Chincholi | B | M | 1 | | | | | 1 |
| 141 | 364 | Morachi Chincholi | B | M | 1 | | | | | 1 |
| 142 | 671 | Morachi Chincholi | B | M | 1 | | | | | 1 |
| 143 | 672 | Morachi Chincholi | B | M | 1 | | | | | 1 |
| 144 | 673 | Morachi Chincholi | B | M | 1 | | | | | 1 |
| 145 | 674 | Morachi Chincholi | B | M | | | 1 | | | 1 |
| 146 | 82 | Morachi Chincholi | B | M | 1 | 1 | | | | 2 |
| 147 | 100 | Morachi Chincholi | B | M | 1 | 1 | | | | 2 |
| 148 | 101 | Morachi Chincholi | B | M | | | 2 | | | 2 |
| 149 | 105 | Morachi Chincholi | B | M | 2 | | | | | 2 |
| 150 | 128 | Morachi Chincholi | B | M | 1 | 1 | | | | 2 |
| 151 | 257 | Morachi Chincholi | B | M | 2 | | | | | 2 |
| 152 | 261 | Morachi Chincholi | B | M | 2 | | | | | 2 |
| 153 | 292 | Morachi Chincholi | B | M | | | 2 | | | 2 |
| 154 | 298 | Morachi Chincholi | B | M | 2 | | | | | 2 |
| 155 | 300 | Morachi Chincholi | B | M | | | 1 | 1 | | 2 |
| 156 | 303 | Morachi Chincholi | B | M | 1 | | 1 | | | 2 |
| 157 | 314 | Morachi Chincholi | B | M | | | 2 | | | 2 |
| 158 | 319 | Morachi Chincholi | B | M | | | 2 | | | 2 |
| 159 | 323 | Morachi Chincholi | B | M | 2 | | | | | 2 |
| 160 | 351 | Morachi Chincholi | B | M | | | 2 | | | 2 |

| | | | | | | | | | | |
|---|---|---|---|---|---|---|---|---|---|---|
| 161 | 358 | Morachi Chincholi | B | M | | | 2 | | | 2 |
| 162 | 532 | Morachi Chincholi | B | M | | 2 | | | | 2 |
| 163 | 632 | Morachi Chincholi | B | M | | 1 | 1 | | | 2 |
| 164 | 645 | Morachi Chincholi | B | M | | 1 | 1 | | | 2 |
| 165 | 651 | Morachi Chincholi | B | M | | | 1 | 1 | | 2 |
| 166 | 656 | Morachi Chincholi | B | M | | | 2 | | | 2 |
| 167 | 657 | Morachi Chincholi | B | M | | | 2 | | | 2 |
| 168 | 665 | Morachi Chincholi | B | M | | 1 | 1 | | | 2 |
| 169 | 305 | Morachi Chincholi | B | M | | 1 | 1 | | | 2 |
| 170 | 329 | Morachi Chincholi | B | M | 2 | | | | | 2 |
| 171 | 541 | Morachi Chincholi | B | M | 1 | 1 | | | | 2 |
| 172 | 84 | Morachi Chincholi | B | M | | | 2 | | | 2 |
| 173 | 630 | Morachi Chincholi | B | M | | | 2 | | | 2 |
| 174 | 637 | Morachi Chincholi | B | M | | | 2 | | | 2 |
| 175 | 343 | Morachi Chincholi | B | M | | 1 | 1 | | | 2 |
| 176 | 70 | Morachi Chincholi | B | M | | 2 | 1 | | | 3 |
| 177 | 123 | Morachi Chincholi | B | M | | 2 | 1 | | | 3 |
| 178 | 294 | Morachi Chincholi | B | M | | | 3 | | | 3 |
| 179 | 325 | Morachi Chincholi | B | M | | | 3 | | | 3 |
| 180 | 336 | Morachi Chincholi | B | M | | | 3 | | | 3 |
| 181 | 350 | Morachi Chincholi | B | M | | 1 | 2 | | | 3 |
| 182 | 368 | Morachi Chincholi | B | M | | | 3 | | | 3 |
| 183 | 530 | Morachi Chincholi | B | M | | | 3 | | | 3 |
| 184 | 534 | Morachi Chincholi | B | M | | | 3 | | | 3 |
| 185 | 644 | Morachi Chincholi | B | M | | 1 | 2 | | | 3 |
| 186 | 690 | Morachi Chincholi | B | M | | 1 | 2 | | | 3 |
| 187 | 536 | Morachi Chincholi | B | M | | 1 | 2 | | | 3 |

| | | | | | | | | | | |
|---|---|---|---|---|---|---|---|---|---|---|
| 188 | 378 | Morachi Chincholi | B | M | | 1 | 2 | | | 3 |
| 189 | 379 | Morachi Chincholi | B | M | | 1 | 2 | | | 3 |
| 190 | 106 | Morachi Chincholi | B | M | | 1 | 2 | | | 3 |
| 191 | 355 | Morachi Chincholi | B | M | | 1 | 2 | | | 3 |
| 192 | 623 | Morachi Chincholi | B | M | | 1 | 2 | | | 3 |
| 193 | 540 | Morachi Chincholi | B | M | 2 | 1 | | | | 3 |
| 194 | 107 | Morachi Chincholi | B | M | | 3 | | | | 3 |
| 195 | 88 | Morachi Chincholi | B | M | 1 | 1 | 2 | | | 4 |
| 196 | 89 | Morachi Chincholi | B | M | | 3 | 1 | | | 4 |
| 197 | 297 | Morachi Chincholi | B | M | | | 4 | | | 4 |
| 198 | 338 | Morachi Chincholi | B | M | | | | | 4 | 4 |
| 199 | 661 | Morachi Chincholi | B | M | | 1 | 3 | | | 4 |
| 200 | 662 | Morachi Chincholi | B | M | | 2 | 2 | | | 4 |
| 201 | 676 | Morachi Chincholi | B | M | 3 | | 1 | | | 4 |
| 202 | 535 | Morachi Chincholi | B | M | | | 4 | | | 4 |
| 203 | 326 | Morachi Chincholi | B | M | | 2 | 2 | | | 4 |
| 204 | 125 | Morachi Chincholi | B | M | 2 | 2 | 1 | | | 5 |
| 205 | 293 | Morachi Chincholi | B | M | | 1 | 3 | 1 | | 5 |
| 206 | 310 | Morachi Chincholi | B | M | | 2 | 3 | | | 5 |
| 207 | 352 | Morachi Chincholi | B | M | | 2 | 3 | | | 5 |
| 208 | 359 | Morachi Chincholi | B | M | | 2 | 3 | | | 5 |
| 209 | 122 | Morachi Chincholi | B | M | | 3 | 2 | 1 | | 6 |
| 210 | 533 | Morachi Chincholi | B | M | | 2 | 4 | | | 6 |
| 211 | 542 | Morachi Chincholi | B | M | | 1 | 4 | | 1 | 6 |
| 212 | 304 | Morachi Chincholi | B | M | | 2 | 4 | | | 6 |
| 213 | 666 | Morachi Chincholi | B | M | | 3 | 4 | | | 7 |
| 214 | 694 | Morachi Chincholi | B | M | | | 2 | 5 | | 7 |

| | | | | | | | | | | |
|---|---|---|---|---|---|---|---|---|---|---|
| 215 | 376 | Morachi Chincholi | B | M | | 6 | 2 | | | 8 |
| 216 | 9 | Morachi Chincholi | NB | PostM | 1 | | | | | 1 |
| 217 | 11 | Morachi Chincholi | NB | PostM | 1 | | | | | 1 |
| 218 | 383 | Morachi Chincholi | NB | PostM | 1 | | | | | 1 |
| 219 | 385 | Morachi Chincholi | NB | PostM | 1 | | | | | 1 |
| 220 | 386 | Morachi Chincholi | NB | PostM | 1 | | | | | 1 |
| 221 | 394 | Morachi Chincholi | NB | PostM | 1 | | | | | 1 |
| 222 | 396 | Morachi Chincholi | NB | PostM | 1 | | | | | 1 |
| 223 | 695 | Morachi Chincholi | NB | PostM | 1 | | | | | 1 |
| 224 | 696 | Morachi Chincholi | NB | PostM | 1 | | | | | 1 |
| 225 | 697 | Morachi Chincholi | NB | PostM | 1 | | | | | 1 |
| 226 | 698 | Morachi Chincholi | NB | PostM | 1 | | | | | 1 |
| 227 | 699 | Morachi Chincholi | NB | PostM | 1 | | | | | 1 |
| 228 | 700 | Morachi Chincholi | NB | PostM | 1 | | | | | 1 |
| 229 | 393 | Morachi Chincholi | NB | PostM | 1 | | | | | 1 |
| 230 | 387 | Morachi Chincholi | NB | PostM | 1 | | | | | 1 |
| 231 | 388 | Morachi Chincholi | NB | PostM | | | 1 | | | 1 |
| 232 | 390 | Morachi Chincholi | NB | PostM | 1 | | | | | 1 |
| 233 | 701 | Morachi Chincholi | NB | PostM | 1 | | | | | 1 |
| 234 | 706 | Morachi Chincholi | NB | PostM | | | 1 | | | 1 |
| 235 | 707 | Morachi Chincholi | NB | PostM | 1 | | | | | 1 |
| 236 | 10 | Morachi Chincholi | NB | PostM | 2 | | | | | 2 |
| 237 | 382 | Morachi Chincholi | NB | PostM | | | 2 | | | 2 |
| 238 | 391 | Morachi Chincholi | NB | PostM | 2 | | | | | 2 |
| 239 | 702 | Morachi Chincholi | NB | PostM | 2 | | | | | 2 |
| 240 | 704 | Morachi Chincholi | NB | PostM | 2 | | | | | 2 |
| 241 | 705 | Morachi Chincholi | NB | PostM | | | 2 | | | 2 |

| | | | | | | | | | | |
|---|---|---|---|---|---|---|---|---|---|---|
| 242 | 6 | Morachi Chincholi | NB | PostM | 1 | 2 | | | | 3 |
| 243 | 131 | Morachi Chincholi | NB | PostM | | | 1 | 2 | | 3 |
| 244 | 395 | Morachi Chincholi | NB | PostM | 1 | | 2 | | | 3 |
| 245 | 389 | Morachi Chincholi | NB | PostM | 1 | 2 | | | | 3 |
| 246 | 703 | Morachi Chincholi | NB | PostM | 2 | | 1 | | | 3 |
| 247 | 7 | Morachi Chincholi | NB | PostM | | | | 1 | 3 | 4 |
| 248 | 392 | Morachi Chincholi | NB | PostM | 2 | 2 | | | | 4 |
| 249 | 5 | Morachi Chincholi | NB | PostM | | 2 | 3 | | | 5 |
| 250 | 8 | Morachi Chincholi | NB | PostM | | | 1 | 5 | | 6 |
| 251 | 384 | Morachi Chincholi | NB | PostM | | 2 | 2 | | 3 | 7 |
| 252 | 12 | Morachi Chincholi | NB | PostM | | | 3 | 6 | | 9 |
| 253 | 139 | Morachi Chincholi | B | PreM | 1 | | | | | 0 |
| 254 | 3 | Morachi Chincholi | B | PreM | 1 | | | | | 1 |
| 255 | 15 | Morachi Chincholi | NB | PreM | 1 | | | | | 1 |
| 256 | 16 | Morachi Chincholi | NB | PreM | 1 | | | | | 1 |
| 257 | 48 | Morachi Chincholi | B | PreM | 1 | | | | | 1 |
| 258 | 49 | Morachi Chincholi | B | PreM | 1 | | | | | 1 |
| 259 | 51 | Morachi Chincholi | B | PreM | 1 | | | | | 1 |
| 260 | 52 | Morachi Chincholi | B | PreM | 1 | | | | | 1 |
| 261 | 53 | Morachi Chincholi | B | PreM | 1 | | | | | 1 |
| 262 | 56 | Morachi Chincholi | B | PreM | 1 | | | | | 1 |
| 263 | 57 | Morachi Chincholi | B | PreM | 1 | | | | | 1 |
| 264 | 60 | Morachi Chincholi | B | PreM | 1 | | | | | 1 |
| 265 | 61 | Morachi Chincholi | B | PreM | 1 | | | | | 1 |
| 266 | 62 | Morachi Chincholi | B | PreM | 1 | | | | | 1 |
| 267 | 133 | Morachi Chincholi | B | PreM | 1 | 0 | 0 | 0 | 0 | 1 |
| 268 | 134 | Morachi Chincholi | B | PreM | 1 | 0 | 0 | 0 | 0 | 1 |

| | | | | | | | | | | |
|---|---|---|---|---|---|---|---|---|---|---|
| 269 | 135 | Morachi Chincholi | B | PreM | 1 | | | | | 1 |
| 270 | 136 | Morachi Chincholi | B | PreM | 1 | | | | | 1 |
| 271 | 140 | Morachi Chincholi | B | PreM | 1 | | | | | 1 |
| 272 | 142 | Morachi Chincholi | B | PreM | 1 | 0 | 0 | 0 | 0 | 1 |
| 273 | 144 | Morachi Chincholi | B | PreM | 1 | | | | | 1 |
| 274 | 145 | Morachi Chincholi | B | PreM | 1 | | | | | 1 |
| 275 | 153 | Morachi Chincholi | B | PreM | 1 | | | | | 1 |
| 276 | 154 | Morachi Chincholi | B | PreM | | | 1 | | | 1 |
| 277 | 155 | Morachi Chincholi | B | PreM | | | 1 | | | 1 |
| 278 | 156 | Morachi Chincholi | B | PreM | | | 1 | | | 1 |
| 279 | 157 | Morachi Chincholi | B | PreM | 1 | | | | | 1 |
| 280 | 161 | Morachi Chincholi | B | PreM | | | | | | 1 |
| 281 | 166 | Morachi Chincholi | B | PreM | | | 1 | | | 1 |
| 282 | 168 | Morachi Chincholi | B | PreM | 1 | | | | | 1 |
| 283 | 169 | Morachi Chincholi | B | PreM | | 1 | | | | 1 |
| 284 | 170 | Morachi Chincholi | B | PreM | 1 | | | | | 1 |
| 285 | 171 | Morachi Chincholi | B | PreM | 1 | | | | | 1 |
| 286 | 172 | Morachi Chincholi | B | PreM | 1 | | | | | 1 |
| 287 | 174 | Morachi Chincholi | B | PreM | | 1 | | | | 1 |
| 288 | 179 | Morachi Chincholi | B | PreM | 1 | | | | | 1 |
| 289 | 180 | Morachi Chincholi | B | PreM | 1 | | | | | 1 |
| 290 | 181 | Morachi Chincholi | B | PreM | 1 | | | | | 1 |
| 291 | 184 | Morachi Chincholi | B | PreM | 1 | | | | | 1 |
| 292 | 195 | Morachi Chincholi | B | PreM | 1 | | | | | 1 |
| 293 | 196 | Morachi Chincholi | B | PreM | 1 | | | | | 1 |
| 294 | 197 | Morachi Chincholi | B | PreM | 1 | | | | | 1 |
| 295 | 198 | Morachi Chincholi | B | PreM | 1 | | | | | 1 |

| | | | | | | | | | | |
|---|---|---|---|---|---|---|---|---|---|---|
| 296 | 200 | Morachi Chincholi | B | PreM | 1 | | | | | 1 |
| 297 | 201 | Morachi Chincholi | B | PreM | | | 1 | | | 1 |
| 298 | 202 | Morachi Chincholi | B | PreM | 1 | | | | | 1 |
| 299 | 203 | Morachi Chincholi | B | PreM | 1 | | | | | 1 |
| 300 | 206 | Morachi Chincholi | B | PreM | 1 | | | | | 1 |
| 301 | 207 | Morachi Chincholi | B | PreM | 1 | | | | | 1 |
| 302 | 208 | Morachi Chincholi | B | PreM | 1 | | | | | 1 |
| 303 | 210 | Morachi Chincholi | B | PreM | | 1 | | | | 1 |
| 304 | 211 | Morachi Chincholi | B | PreM | 1 | | | | | 1 |
| 305 | 212 | Morachi Chincholi | B | PreM | | 1 | | | | 1 |
| 306 | 215 | Morachi Chincholi | B | PreM | | 1 | | | | 1 |
| 307 | 216 | Morachi Chincholi | B | PreM | 1 | | | | | 1 |
| 308 | 219 | Morachi Chincholi | B | PreM | 1 | | | | | 1 |
| 309 | 222 | Morachi Chincholi | B | PreM | 1 | | | | | 1 |
| 310 | 223 | Morachi Chincholi | B | PreM | 1 | | | | | 1 |
| 311 | 224 | Morachi Chincholi | B | PreM | | | 1 | | | 1 |
| 312 | 225 | Morachi Chincholi | B | PreM | 1 | | | | | 1 |
| 313 | 226 | Morachi Chincholi | B | PreM | | 1 | | | | 1 |
| 314 | 227 | Morachi Chincholi | B | PreM | | 1 | | | | 1 |
| 315 | 230 | Morachi Chincholi | B | PreM | 1 | | | | | 1 |
| 316 | 423 | Morachi Chincholi | NB | PreM | 1 | | | | | 1 |
| 317 | 425 | Morachi Chincholi | B | PreM | 1 | | | | | 1 |
| 318 | 426 | Morachi Chincholi | B | PreM | 1 | | | | | 1 |
| 319 | 434 | Morachi Chincholi | B | PreM | 1 | | | | | 1 |
| 320 | 435 | Morachi Chincholi | B | PreM | 1 | | | | | 1 |
| 321 | 437 | Morachi Chincholi | B | PreM | 1 | | | | | 1 |
| 322 | 441 | Morachi Chincholi | B | PreM | | | 1 | | | 1 |

| | | | | | | | | | | |
|---|---|---|---|---|---|---|---|---|---|---|
| 323 | 442 | Morachi Chincholi | B | PreM | 1 | | | | | 1 |
| 324 | 444 | Morachi Chincholi | B | PreM | 1 | | | | | 1 |
| 325 | 448 | Morachi Chincholi | B | PreM | | | 1 | | | 1 |
| 326 | 451 | Morachi Chincholi | B | PreM | 1 | | | | | 1 |
| 327 | 452 | Morachi Chincholi | B | PreM | 1 | | | | | 1 |
| 328 | 453 | Morachi Chincholi | B | PreM | 1 | | | | | 1 |
| 329 | 454 | Morachi Chincholi | B | PreM | 1 | | | | | 1 |
| 330 | 464 | Morachi Chincholi | B | PreM | 1 | | | | | 1 |
| 331 | 501 | Morachi Chincholi | B | PreM | 1 | | | | | 1 |
| 332 | 503 | Morachi Chincholi | B | PreM | 1 | | | | | 1 |
| 333 | 504 | Morachi Chincholi | B | PreM | 1 | | | | | 1 |
| 334 | 505 | Morachi Chincholi | B | PreM | | | 1 | | | 1 |
| 335 | 506 | Morachi Chincholi | B | PreM | 1 | | | | | 1 |
| 336 | 508 | Morachi Chincholi | B | PreM | 1 | | | | | 1 |
| 337 | 509 | Morachi Chincholi | B | PreM | 1 | | | | | 1 |
| 338 | 510 | Morachi Chincholi | B | PreM | 1 | | | | | 1 |
| 339 | 514 | Morachi Chincholi | B | PreM | 1 | | | | | 1 |
| 340 | 517 | Morachi Chincholi | B | PreM | 1 | | | | | 1 |
| 341 | 521 | Morachi Chincholi | B | PreM | 1 | | | | | 1 |
| 342 | 524 | Morachi Chincholi | B | PreM | | | 1 | | | 1 |
| 343 | 525 | Morachi Chincholi | B | PreM | | 1 | | | | 1 |
| 344 | 526 | Morachi Chincholi | B | PreM | | 1 | | | | 1 |
| 345 | 138 | Morachi Chincholi | B | PreM | 1 | | | | | 1 |
| 346 | 17 | Morachi Chincholi | NB | PreM | 1 | | | | | 1 |
| 347 | 446 | Morachi Chincholi | B | PreM | 1 | | | | | 1 |
| 348 | 406 | Morachi Chincholi | NB | PreM | 1 | | | | | 1 |
| 349 | 449 | Morachi Chincholi | B | PreM | 1 | | | | | 1 |

| | | | | | | | | | | |
|---|---|---|---|---|---|---|---|---|---|---|
| 350 | 459 | Morachi Chincholi | B | PreM | 1 | | | | | 1 |
| 351 | 460 | Morachi Chincholi | B | PreM | 1 | | | | | 1 |
| 352 | 173 | Morachi Chincholi | B | PreM | 1 | | | | | 1 |
| 353 | 228 | Morachi Chincholi | B | PreM | 1 | | | | | 1 |
| 354 | 229 | Morachi Chincholi | B | PreM | 1 | | | | | 1 |
| 355 | 232 | Morachi Chincholi | B | PreM | | | 1 | | | 1 |
| 356 | 233 | Morachi Chincholi | B | PreM | 1 | | | | | 1 |
| 357 | 234 | Morachi Chincholi | B | PreM | 1 | | | | | 1 |
| 358 | 238 | Morachi Chincholi | B | PreM | 1 | | | | | 1 |
| 359 | 447 | Morachi Chincholi | B | PreM | 1 | | | | | 1 |
| 360 | 235 | Morachi Chincholi | B | PreM | 1 | | | | | 1 |
| 361 | 236 | Morachi Chincholi | B | PreM | | 1 | | | | 1 |
| 362 | 237 | Morachi Chincholi | B | PreM | 1 | | | | | 1 |
| 363 | 411 | Morachi Chincholi | NB | PreM | 1 | | | | | 1 |
| 364 | 412 | Morachi Chincholi | NB | PreM | 1 | | | | | 1 |
| 365 | 413 | Morachi Chincholi | NB | PreM | | | 1 | | | 1 |
| 366 | 414 | Morachi Chincholi | NB | PreM | 1 | | | | | 1 |
| 367 | 415 | Morachi Chincholi | NB | PreM | 1 | | | | | 1 |
| 368 | 416 | Morachi Chincholi | NB | PreM | 1 | | | | | 1 |
| 369 | 417 | Morachi Chincholi | NB | PreM | 1 | | | | | 1 |
| 370 | 418 | Morachi Chincholi | NB | PreM | 1 | | | | | 1 |
| 371 | 420 | Morachi Chincholi | NB | PreM | 1 | | | | | 1 |
| 372 | 429 | Morachi Chincholi | B | PreM | 1 | | | | | 1 |
| 373 | 430 | Morachi Chincholi | B | PreM | 1 | | | | | 1 |
| 374 | 432 | Morachi Chincholi | B | PreM | 1 | | | | | 1 |
| 375 | 433 | Morachi Chincholi | B | PreM | 1 | | | | | 1 |
| 376 | 148 | Morachi Chincholi | B | PreM | 1 | | | | | 1 |

| | | | | | | | | | | | |
|---|---|---|---|---|---|---|---|---|---|---|---|
| 377 | 149 | Moracchi Chincholi | B | PreM | 1 | | | | | | 1 |
| 378 | 160 | Moracchi Chincholi | B | PreM | 1 | | | | | | 1 |
| 379 | 165 | Moracchi Chincholi | B | PreM | 1 | | | | | | 1 |
| 380 | 205 | Moracchi Chincholi | B | PreM | 1 | | | | | | 1 |
| 381 | 189 | Moracchi Chincholi | B | PreM | 1 | | | | | | 1 |
| 382 | 1 | Moracchi Chincholi | B | PreM | 1 | | | 1 | | | 2 |
| 383 | 50 | Moracchi Chincholi | B | PreM | 2 | | | | | | 2 |
| 384 | 54 | Moracchi Chincholi | B | PreM | | | | 2 | | | 2 |
| 385 | 55 | Moracchi Chincholi | B | PreM | | | 2 | | | | 2 |
| 386 | 59 | Moracchi Chincholi | B | PreM | 2 | | | | | | 2 |
| 387 | 67 | Moracchi Chincholi | B | PreM | 1 | | 1 | | | | 2 |
| 388 | 146 | Moracchi Chincholi | B | PreM | | | 2 | | | | 2 |
| 389 | 152 | Moracchi Chincholi | B | PreM | | 2 | | | | | 2 |
| 390 | 163 | Moracchi Chincholi | B | PreM | 2 | | | | | | 2 |
| 391 | 176 | Moracchi Chincholi | B | PreM | 2 | | | | | | 2 |
| 392 | 178 | Moracchi Chincholi | B | PreM | 2 | | | | | | 2 |
| 393 | 182 | Moracchi Chincholi | B | PreM | | | 2 | | | | 2 |
| 394 | 186 | Moracchi Chincholi | B | PreM | 1 | | | | | | 2 |
| 395 | 191 | Moracchi Chincholi | B | PreM | | 2 | | | | | 2 |
| 396 | 192 | Moracchi Chincholi | B | PreM | | | 1 | 1 | | | 2 |
| 397 | 209 | Moracchi Chincholi | B | PreM | | | 2 | | | | 2 |
| 398 | 440 | Moracchi Chincholi | B | PreM | 2 | | | | | | 2 |
| 399 | 443 | Moracchi Chincholi | B | PreM | | 2 | | | | | 2 |
| 400 | 462 | Moracchi Chincholi | B | PreM | 1 | 1 | | | | | 2 |
| 401 | 463 | Moracchi Chincholi | B | PreM | | 2 | | | | | 2 |
| 402 | 465 | Moracchi Chincholi | B | PreM | | | 2 | | | | 2 |
| 403 | 512 | Moracchi Chincholi | B | PreM | 1 | 1 | | | | | 2 |

| | | | | | | | | | | |
|---|---|---|---|---|---|---|---|---|---|---|
| 404 | 513 | Morachi Chincholi | B | PreM | | | 2 | | | 2 |
| 405 | 519 | Morachi Chincholi | B | PreM | 2 | | | | | 2 |
| 406 | 438 | Morachi Chincholi | B | PreM | 1 | 1 | | | | 2 |
| 407 | 498 | Morachi Chincholi | B | PreM | 1 | 1 | | | | 2 |
| 408 | 499 | Morachi Chincholi | B | PreM | 1 | 1 | | | | 2 |
| 409 | 500 | Morachi Chincholi | B | PreM | 1 | 1 | | | | 2 |
| 410 | 520 | Morachi Chincholi | B | PreM | | | 2 | | | 2 |
| 411 | 461 | Morachi Chincholi | B | PreM | 1 | 1 | | | | 2 |
| 412 | 431 | Morachi Chincholi | B | PreM | | | 2 | | | 2 |
| 413 | 147 | Morachi Chincholi | B | PreM | 0 | 1 | 1 | | | 2 |
| 414 | 158 | Morachi Chincholi | B | PreM | | | 2 | | | 2 |
| 415 | 183 | Morachi Chincholi | B | PreM | 2 | | | | | 2 |
| 416 | 456 | Morachi Chincholi | B | PreM | 2 | | | | | 2 |
| 417 | 2 | Morachi Chincholi | B | PreM | 1 | | 2 | | | 3 |
| 418 | 4 | Morachi Chincholi | B | PreM | 2 | | 1 | | | 3 |
| 419 | 63 | Morachi Chincholi | B | PreM | | | | 3 | | 3 |
| 420 | 65 | Morachi Chincholi | B | PreM | | | 1 | 2 | | 3 |
| 421 | 69 | Morachi Chincholi | B | PreM | 1 | | 2 | | | 3 |
| 422 | 141 | Morachi Chincholi | B | PreM | 0 | 1 | 2 | | | 3 |
| 423 | 193 | Morachi Chincholi | B | PreM | | | 3 | | | 3 |
| 424 | 199 | Morachi Chincholi | B | PreM | | 2 | 1 | | | 3 |
| 425 | 217 | Morachi Chincholi | B | PreM | | | 3 | | | 3 |
| 426 | 231 | Morachi Chincholi | B | PreM | | 1 | 2 | | | 3 |
| 427 | 404 | Morachi Chincholi | NB | PreM | 2 | | 1 | | | 3 |
| 428 | 445 | Morachi Chincholi | B | PreM | 1 | 2 | | | | 3 |
| 429 | 450 | Morachi Chincholi | B | PreM | 1 | | 2 | | | 3 |
| 430 | 455 | Morachi Chincholi | B | PreM | | | 3 | | | 3 |

| | | | | | | | | | | |
|---|---|---|---|---|---|---|---|---|---|---|
| 431 | 507 | Morachi Chincholi | B | PreM | | | 3 | | | 3 |
| 432 | 14 | Morachi Chincholi | NB | PreM | 3 | | | | | 3 |
| 433 | 496 | Morachi Chincholi | B | PreM | 1 | 1 | 1 | | | 3 |
| 434 | 497 | Morachi Chincholi | B | PreM | 1 | 2 | | | | 3 |
| 435 | 457 | Morachi Chincholi | B | PreM | 2 | 1 | | | | 3 |
| 436 | 421 | Morachi Chincholi | NB | PreM | | 1 | 2 | | | 3 |
| 437 | 428 | Morachi Chincholi | B | PreM | | | 3 | | | 3 |
| 438 | 439 | Morachi Chincholi | B | PreM | | 2 | 1 | | | 3 |
| 439 | 132 | Morachi Chincholi | B | PreM | 0 | 0 | 4 | 0 | 0 | 4 |
| 440 | 137 | Morachi Chincholi | B | PreM | 0 | 0 | 4 | | | 4 |
| 441 | 167 | Morachi Chincholi | B | PreM | 1 | | 3 | | | 4 |
| 442 | 190 | Morachi Chincholi | B | PreM | | | 4 | | | 4 |
| 443 | 194 | Morachi Chincholi | B | PreM | | 2 | 2 | | | 4 |
| 444 | 204 | Morachi Chincholi | B | PreM | 1 | 1 | 2 | | | 4 |
| 445 | 220 | Morachi Chincholi | B | PreM | 1 | 1 | 2 | | | 4 |
| 446 | 495 | Morachi Chincholi | B | PreM | | 3 | 1 | | | 4 |
| 447 | 502 | Morachi Chincholi | B | PreM | 1 | | 3 | | | 4 |
| 448 | 511 | Morachi Chincholi | B | PreM | | | 4 | | | 4 |
| 449 | 188 | Morachi Chincholi | B | PreM | | 1 | 3 | | | 4 |
| 450 | 419 | Morachi Chincholi | NB | PreM | | 1 | 3 | | | 4 |
| 451 | 159 | Morachi Chincholi | B | PreM | | 1 | 3 | | | 4 |
| 452 | 164 | Morachi Chincholi | B | PreM | | 1 | 2 | | 1 | 4 |
| 453 | 13 | Morachi Chincholi | NB | PreM | 5 | | | | | 5 |
| 454 | 58 | Morachi Chincholi | B | PreM | 1 | | 4 | | | 5 |
| 455 | 162 | Morachi Chincholi | B | PreM | | 1 | 4 | | | 5 |
| 456 | 177 | Morachi Chincholi | B | PreM | | | 4 | 1 | | 5 |
| 457 | 213 | Morachi Chincholi | B | PreM | 2 | 1 | 2 | | | 5 |

| | | | | | | | | | | |
|---|---|---|---|---|---|---|---|---|---|---|
| 458 | 422 | Morachi Chincholi | NB | PreM | 1 | 2 | 2 | | | 5 |
| 459 | 424 | Morachi Chincholi | NB | PreM | | 2 | 3 | | | 5 |
| 460 | 436 | Morachi Chincholi | B | PreM | | 2 | 3 | | | 5 |
| 461 | 522 | Morachi Chincholi | B | PreM | 1 | 2 | 2 | | | 5 |
| 462 | 523 | Morachi Chincholi | B | PreM | 1 | 2 | 3 | | | 5 |
| 463 | 185 | Morachi Chincholi | B | PreM | | | 1 | 4 | | 5 |
| 464 | 516 | Morachi Chincholi | B | PreM | | 2 | 3 | | | 5 |
| 465 | 458 | Morachi Chincholi | B | PreM | 2 | 2 | 1 | | | 5 |
| 466 | 427 | Morachi Chincholi | B | PreM | | 2 | 3 | | | 5 |
| 467 | 143 | Morachi Chincholi | B | PreM | 0 | 2 | 3 | | | 5 |
| 468 | 28 | Morachi Chincholi | NB | PreM | | | | | 6 | 6 |
| 469 | 66 | Morachi Chincholi | B | PreM | 1 | 1 | 4 | | | 6 |
| 470 | 218 | Morachi Chincholi | B | PreM | 2 | 3 | 1 | | | 6 |
| 471 | 187 | Morachi Chincholi | B | PreM | | 2 | 4 | | | 6 |
| 472 | 466 | Morachi Chincholi | B | PreM | | 2 | 4 | | | 6 |
| 473 | 515 | Morachi Chincholi | B | PreM | 1 | 4 | 1 | | | 6 |
| 474 | 405 | Morachi Chincholi | NB | PreM | | | 6 | | | 6 |
| 475 | 27 | Morachi Chincholi | NB | PreM | 1 | 1 | 5 | | | 7 |
| 476 | 150 | Morachi Chincholi | B | PreM | 1 | 1 | 5 | | | 7 |
| 477 | 151 | Morachi Chincholi | B | PreM | | 1 | 6 | | | 7 |
| 478 | 214 | Morachi Chincholi | B | PreM | | 5 | 2 | | | 7 |
| 479 | 221 | Morachi Chincholi | B | PreM | 2 | 2 | 3 | | | 7 |
| 480 | 64 | Morachi Chincholi | B | PreM | | 3 | 5 | | | 8 |
| 481 | 175 | Morachi Chincholi | B | PreM | 5 | | 3 | | | 8 |
| 482 | 518 | Morachi Chincholi | B | PreM | | 3 | 5 | | | 8 |
| 483 | 68 | Morachi Chincholi | B | PreM | | 2 | 6 | | | 8 |
| 484 | 29 | Morachi Chincholi | NB | PreM | 1 | 1 | 10 | | | 12 |

| | | | | | | | | | | |
|---|---|---|---|---|---|---|---|---|---|---|
| 485 | 277 | Nashik | B | M | 1 | | | | | 1 |
| 486 | 279 | Nashik | B | M | | | 1 | | | 1 |
| 487 | 281 | Nashik | B | M | | 1 | | | | 1 |
| 488 | 109 | Nashik | B | M | 1 | | | | | 1 |
| 489 | 111 | Nashik | B | M | 1 | | | | | 1 |
| 490 | 112 | Nashik | B | M | 1 | | | | | 1 |
| 491 | 113 | Nashik | B | M | 1 | | | | | 1 |
| 492 | 114 | Nashik | B | M | | | 1 | | | 1 |
| 493 | 115 | Nashik | B | M | 1 | | | | | 1 |
| 494 | 118 | Nashik | B | M | | | 1 | | | 1 |
| 495 | 119 | Nashik | B | M | | | | 1 | | 1 |
| 496 | 120 | Nashik | B | M | 1 | | | | | 1 |
| 497 | 262 | Nashik | B | M | | | 1 | | | 1 |
| 498 | 263 | Nashik | B | M | 1 | | | | | 1 |
| 499 | 264 | Nashik | B | M | | | 1 | | | 1 |
| 500 | 265 | Nashik | B | M | 1 | | | | | 1 |
| 501 | 266 | Nashik | B | M | 1 | | | | | 1 |
| 502 | 267 | Nashik | B | M | 1 | | | | | 1 |
| 503 | 268 | Nashik | B | M | 1 | | | | | 1 |
| 504 | 269 | Nashik | B | M | 1 | | | | | 1 |
| 505 | 270 | Nashik | B | M | 1 | | | | | 1 |
| 506 | 271 | Nashik | B | M | 1 | | | | | 1 |
| 507 | 272 | Nashik | B | M | 1 | | | | | 1 |
| 508 | 273 | Nashik | B | M | 1 | | | | | 1 |
| 509 | 275 | Nashik | B | M | 1 | | | | | 1 |
| 510 | 284 | Nashik | B | M | 1 | | | | | 1 |
| 511 | 286 | Nashik | B | M | 1 | | | | | 1 |
| 512 | 274 | Nashik | B | M | | | 2 | | | 2 |
| 513 | 116 | Nashik | B | M | 2 | | | | | 2 |
| 514 | 117 | Nashik | B | M | 1 | 1 | | | | 2 |
| 515 | 285 | Nashik | B | M | 1 | | 1 | | | 2 |
| 516 | 276 | Nashik | B | M | | 2 | 1 | | | 3 |
| 517 | 280 | Nashik | B | M | | | 3 | | | 3 |
| 518 | 283 | Nashik | B | M | | 2 | 1 | | | 3 |
| 519 | 278 | Nashik | B | M | | 2 | 2 | | | 4 |
| 520 | 121 | Nashik | B | M | 1 | 1 | 2 | | | 4 |
| 521 | 110 | Nashik | B | M | | 2 | 1 | 2 | | 5 |
| 522 | 282 | Nashik | B | M | | | 6 | | | 6 |
| 523 | 748 | Nashik | NB | PostM | 1 | | | | | 1 |
| 524 | 749 | Nashik | NB | PostM | 1 | | | | | 1 |
| 525 | 750 | Nashik | NB | PostM | 1 | | | | | 1 |
| 526 | 751 | Nashik | NB | PostM | 1 | | | | | 1 |
| 527 | 752 | Nashik | NB | PostM | 1 | | | | | 1 |
| 528 | 753 | Nashik | NB | PostM | 1 | | | | | 1 |
| 529 | 754 | Nashik | NB | PostM | 1 | | | | | 1 |

| | | | | | | | | | | |
|---|---|---|---|---|---|---|---|---|---|---|
| 530 | 756 | Nashik | NB | PostM | 1 | | | | | 1 |
| 531 | 757 | Nashik | NB | PostM | | | 1 | | | 1 |
| 532 | 758 | Nashik | NB | PostM | | | 1 | | | 1 |
| 533 | 759 | Nashik | NB | PostM | | | 1 | | | 1 |
| 534 | 764 | Nashik | NB | PostM | | | 1 | | | 1 |
| 535 | 768 | Nashik | NB | PostM | 1 | | | | | 1 |
| 536 | 769 | Nashik | NB | PostM | 1 | | | | | 1 |
| 537 | 130 | Nashik | NB | PostM | 2 | | | | | 2 |
| 538 | 760 | Nashik | NB | PostM | 2 | | | | | 2 |
| 539 | 762 | Nashik | NB | PostM | 2 | | | | | 2 |
| 540 | 765 | Nashik | NB | PostM | 2 | | | | | 2 |
| 541 | 766 | Nashik | NB | PostM | 2 | | | | | 2 |
| 542 | 770 | Nashik | NB | PostM | | | 2 | | | 2 |
| 543 | 755 | Nashik | NB | PostM | 3 | | | | | 3 |
| 544 | 767 | Nashik | NB | PostM | | | 2 | 1 | | 3 |
| 545 | 761 | Nashik | NB | PostM | 1 | | 1 | 2 | | 4 |
| 546 | 763 | Nashik | NB | PostM | | | 1 | 3 | | 4 |
| 547 | 471 | Nashik | B | PreM | 1 | | | | | 1 |
| 548 | 32 | Nashik | B | PreM | 1 | | | | | 1 |
| 549 | 37 | Nashik | B | PreM | 1 | | | | | 1 |
| 550 | 41 | Nashik | B | PreM | 1 | | | | | 1 |
| 551 | 45 | Nashik | B | PreM | 1 | | | | | 1 |
| 552 | 467 | Nashik | B | PreM | | 1 | | | | 1 |
| 553 | 469 | Nashik | B | PreM | 1 | | | | | 1 |
| 554 | 473 | Nashik | B | PreM | | | 1 | | | 1 |
| 555 | 474 | Nashik | B | PreM | 1 | | | | | 1 |
| 556 | 475 | Nashik | B | PreM | 1 | | | | | 1 |
| 557 | 477 | Nashik | B | PreM | 1 | | | | | 1 |
| 558 | 479 | Nashik | B | PreM | 1 | | | | | 1 |
| 559 | 480 | Nashik | B | PreM | 1 | | | | | 1 |
| 560 | 481 | Nashik | B | PreM | 1 | | | | | 1 |
| 561 | 484 | Nashik | B | PreM | 1 | | | | | 1 |
| 562 | 485 | Nashik | B | PreM | 1 | | | | | 1 |
| 563 | 486 | Nashik | B | PreM | 1 | | | | | 1 |
| 564 | 487 | Nashik | B | PreM | 1 | | | | | 1 |
| 565 | 489 | Nashik | B | PreM | 1 | | | | | 1 |
| 566 | 491 | Nashik | B | PreM | 1 | | | | | 1 |
| 567 | 492 | Nashik | B | PreM | | | 1 | | | 1 |
| 568 | 493 | Nashik | B | PreM | 1 | | | | | 1 |
| 569 | 494 | Nashik | B | PreM | 1 | | | | | 1 |
| 570 | 482 | Nashik | B | PreM | 1 | | 1 | | | 2 |
| 571 | 35 | Nashik | B | PreM | 2 | | | | | 2 |
| 572 | 36 | Nashik | B | PreM | 2 | | | | | 2 |
| 573 | 38 | Nashik | B | PreM | 2 | | | | | 2 |
| 574 | 43 | Nashik | B | PreM | 2 | | | | | 2 |

| | | | | | | | | | | |
|---|---|---|---|---|---|---|---|---|---|---|
| 575 | 44 | Nashik | B | PreM | 1 | | 1 | | | 2 |
| 576 | 46 | Nashik | B | PreM | 3 | | | | | 3 |
| 577 | 470 | Nashik | B | PreM | | 2 | 1 | | | 3 |
| 578 | 483 | Nashik | B | PreM | | 3 | | | | 3 |
| 579 | 31 | Nashik | B | PreM | 3 | | | | | 3 |
| 580 | 33 | Nashik | B | PreM | 2 | | 1 | | | 3 |
| 581 | 39 | Nashik | B | PreM | 3 | | | | | 3 |
| 582 | 468 | Nashik | B | PreM | | | 3 | | | 3 |
| 583 | 478 | Nashik | B | PreM | | | 3 | | | 3 |
| 584 | 488 | Nashik | B | PreM | | 1 | 2 | | | 3 |
| 585 | 490 | Nashik | B | PreM | 3 | | | | | 3 |
| 586 | 40 | Nashik | B | PreM | 1 | | 3 | | | 4 |
| 587 | 30 | Nashik | B | PreM | 1 | 1 | 2 | | | 4 |
| 588 | 47 | Nashik | B | PreM | 2 | 1 | 2 | | | 5 |
| 589 | 34 | Nashik | B | PreM | 1 | | 4 | | | 5 |
| 590 | 42 | Nashik | B | PreM | | | 7 | | | 7 |
| 591 | 472 | Nashik | B | PreM | | 3 | 5 | | | 8 |
| 592 | 476 | Nashik | B | PreM | | 5 | 4 | | | 9 |
| 593 | 594 | Rajasthan | B | M | 1 | | | | | 1 |
| 594 | 596 | Rajasthan | B | M | 1 | | | | | 1 |
| 595 | 597 | Rajasthan | B | M | 1 | | | | | 1 |
| 596 | 598 | Rajasthan | B | M | | | 1 | | | 1 |
| 597 | 599 | Rajasthan | B | M | 1 | | | | | 1 |
| 598 | 600 | Rajasthan | B | M | 1 | | | | | 1 |
| 599 | 601 | Rajasthan | B | M | | 1 | | | | 1 |
| 600 | 602 | Rajasthan | B | M | 1 | | | | | 1 |
| 601 | 604 | Rajasthan | B | M | | | 1 | | | 1 |
| 602 | 605 | Rajasthan | B | M | 1 | | | | | 1 |
| 603 | 606 | Rajasthan | B | M | 1 | | | | | 1 |
| 604 | 608 | Rajasthan | B | M | 1 | | | | | 1 |
| 605 | 612 | Rajasthan | B | M | | 1 | | | | 1 |
| 606 | 613 | Rajasthan | B | M | | | 1 | | | 1 |
| 607 | 616 | Rajasthan | B | M | | | 1 | | | 1 |
| 608 | 617 | Rajasthan | B | M | 1 | | | | | 1 |
| 609 | 622 | Rajasthan | B | M | 1 | | | | | 1 |
| 610 | 595 | Rajasthan | B | M | | | 2 | | | 2 |
| 611 | 603 | Rajasthan | B | M | 2 | | | | | 2 |
| 612 | 609 | Rajasthan | B | M | | 2 | | | | 2 |
| 613 | 611 | Rajasthan | B | M | 1 | 1 | | | | 2 |
| 614 | 620 | Rajasthan | B | M | | | 2 | | | 2 |
| 615 | 621 | Rajasthan | B | M | 1 | 1 | | | | 2 |
| 616 | 619 | Rajasthan | B | M | | 2 | 2 | | | 4 |
| 617 | 610 | Rajasthan | B | M | | 3 | 2 | | | 5 |
| 618 | 618 | Rajasthan | B | M | | 3 | 2 | | | 5 |
| 619 | 620 | Rajasthan | B | M | | 1 | 4 | | | 5 |

| | | | | | | | | | | |
|---|---|---|---|---|---|---|---|---|---|---|
| 620 | 614 | Rajasthan | B | M | | 3 | 3 | | | 6 |
| 621 | 615 | Rajasthan | B | M | | | 6 | | | 6 |
| 622 | 607 | Rajasthan | B | M | | 4 | 3 | | | 7 |
| 623 | 400 | Rajasthan | NB | PostM | 1 | | | | | 1 |
| 624 | 402 | Rajasthan | NB | PostM | 1 | | | | | 1 |
| 625 | 403 | Rajasthan | NB | PostM | 1 | | | | | 1 |
| 626 | 398 | Rajasthan | NB | PostM | 1 | | | | | 1 |
| 627 | 399 | Rajasthan | NB | PostM | | | 1 | | | 1 |
| 628 | 397 | Rajasthan | NB | PostM | 1 | | | | | 1 |
| 629 | 401 | Rajasthan | NB | PostM | 3 | | | | | 3 |
| 630 | 73 | Rajasthan | B | PreM | 1 | | | | | 1 |
| 631 | 239 | Rajasthan | B | PreM | 1 | | | | | 1 |
| 632 | 241 | Rajasthan | B | PreM | 1 | | | | | 1 |
| 633 | 246 | Rajasthan | B | PreM | 1 | | | | | 1 |
| 634 | 247 | Rajasthan | B | PreM | 1 | | | | | 1 |
| 635 | 71 | Rajasthan | B | PreM | 1 | | | | | 1 |
| 636 | 72 | Rajasthan | B | PreM | 1 | | | | | 1 |
| 637 | 74 | Rajasthan | B | PreM | | | 1 | | | 1 |
| 638 | 410 | Rajasthan | NB | PreM | 1 | | | | | 1 |
| 639 | 19 | Rajasthan | NB | PreM | 1 | | | | | 1 |
| 640 | 248 | Rajasthan | B | PreM | 1 | | | | | 1 |
| 641 | 254 | Rajasthan | B | PreM | 1 | | | | | 1 |
| 642 | 255 | Rajasthan | B | PreM | | | 1 | | | 1 |
| 643 | 249 | Rajasthan | B | PreM | | 1 | | | | 1 |
| 644 | 75 | Rajasthan | B | PreM | 1 | | | | | 1 |
| 645 | 244 | Rajasthan | B | PreM | 1 | 1 | | | | 2 |
| 646 | 251 | Rajasthan | B | PreM | | 1 | 1 | | | 2 |
| 647 | 407 | Rajasthan | NB | PreM | 2 | | | | | 2 |
| 648 | 20 | Rajasthan | NB | PreM | 2 | | | | | 2 |
| 649 | 409 | Rajasthan | NB | PreM | 2 | | | | | 2 |
| 650 | 18 | Rajasthan | NB | PreM | 1 | | 1 | | | 2 |
| 651 | 245 | Rajasthan | B | PreM | | | 3 | | | 3 |
| 652 | 252 | Rajasthan | B | PreM | | 2 | 1 | | | 3 |
| 653 | 256 | Rajasthan | B | PreM | | 1 | 2 | | | 3 |
| 654 | 25 | Rajasthan | NB | PreM | | 1 | 3 | | | 4 |
| 655 | 23 | Rajasthan | NB | PreM | 1 | 2 | 2 | | | 5 |
| 656 | 240 | Rajasthan | B | PreM | | | 5 | | | 5 |
| 657 | 21 | Rajasthan | NB | PreM | | 2 | | 3 | | 5 |
| 658 | 242 | Rajasthan | B | PreM | | | 4 | 2 | | 6 |
| 659 | 243 | Rajasthan | B | PreM | | 2 | 4 | | | 6 |
| 660 | 250 | Rajasthan | B | PreM | | 2 | 4 | | | 6 |
| 661 | 253 | Rajasthan | B | PreM | | 1 | 5 | | | 6 |
| 662 | 26 | Rajasthan | NB | PreM | | | 6 | | | 6 |
| 663 | 22 | Rajasthan | NB | PreM | | 2 | | 4 | | 6 |
| 664 | 24 | Rajasthan | NB | PreM | 1 | 2 | 3 | | | 6 |

| | | | | | | | | | | |
|---|---|---|---|---|---|---|---|---|---|---|
| 665 | 408 | Rajasthan | NB | PreM | 7 | | 7 | | | 14 |
